# Wiring logic of the early rodent olfactory system revealed by high-throughput sequencing of single neuron projections

**DOI:** 10.1101/2021.05.12.443929

**Authors:** Yushu Chen, Xiaoyin Chen, Batuhan Baserdem, Huiqing Zhan, Yan Li, Martin B. Davis, Justus M. Kebschull, Anthony M. Zador, Alexei A. Koulakov, Dinu F. Albeanu

**Affiliations:** Cold Spring Harbor Laboratory, Cold Spring Harbor, NY, 11724; Stony Brook University, Stony Brook, NY; Department of Biomedical Engineering, Johns Hopkins School of Medicine, Baltimore, MD 21205, USA

## Abstract

The structure of neuronal connectivity often provides insights into the relevant stimulus features, such as spatial location, orientation, sound frequency, etc^1–6^. The olfactory system, however, appears to lack structured connectivity as suggested by reports of broad and distributed connections both from the olfactory bulb to the piriform cortex^7–22^ and within the cortex^23–25^. These studies have inspired computational models of circuit function that rely on random connectivity^26–33^. It remains, nonetheless, unclear whether the olfactory connectivity contains spatial structure. Here, we use high throughput anatomical methods (MAPseq and BARseq)^34–38^ to analyze the projections of 5,309 bulb and 30,433 piriform cortex output neurons in the mouse at single-cell resolution. We identify previously unrecognized spatial organization in connectivity along the anterior-posterior axis (A-P) of the piriform cortex. We find that both the bulb projections to the cortex and the cortical outputs are not random, but rather form gradients along the A-P axis. Strikingly, these gradients are *matched*: bulb neurons targeting a given location within the piriform cortex co-innervate extra-piriform regions that receive strong inputs from neurons within that piriform locus. We also identify signatures of local connectivity in the piriform cortex. Our findings suggest an organizing principle of matched direct and indirect olfactory pathways that innervate extra-piriform targets in a coordinated manner, thus supporting models of information processing that rely on structured connectivity within the olfactory system.

## Main

The structure of intra- and inter-brain region connectivity provides a scaffolding for neural computation, and different computations rely on different statistics of connections^1–6^. Determining the logic of connectivity is therefore critical for understanding the principles underlying neural processing. In the olfactory system, the axons of olfactory sensory neurons expressing a given odorant receptor type converge to the same glomerulus on the olfactory bulb (OB) surface, forming a 2D stereotypical input map^39–41^. Odorants are represented by the pattern of olfactory sensory neurons activity reflected in the activation of corresponding glomeruli. The spatio-temporal pattern of glomerular activation is routed by the principal output neurons of the bulb called mitral (MC) and tufted cells (TC), which draw excitatory input from individual glomeruli and integrate largely inhibitory lateral and top-down feedback signals arriving from other brain structures^40, 42–46^. Over the past decades, bulk labeling^7–10, 47^ and axonal tracing studies, in conjunction with dye labeling of single glomerular outputs^13–16^ have documented broad and unstructured innervation of large synaptic territories within the piriform cortex (PC), the principal projection target of the bulb, by individual mitral and tufted cell axons. This stands in contrast with the precise mapping of glomerular bulb inputs^39–41, 48^ as well as spatially organized representations ubiquitous in other sensory modalities (vision, audition, somato-sensation, etc.^1, 2, 4–6)^. Inspired by these results and by observations of broad connectivity within the piriform cortex^23–25, 49^, many computational models have assumed random connectivity schemes between the olfactory bulb and piriform cortex^26, 30–32^, and distributed and long-range connectivity within piriform cortex^26, 30, 31, 50, 51^ and other brain regions^27–29, 33, 52^. In these models, the random connectivity is a necessary basis for learning arbitrary stimulus combinations during an individual’s lifetime. It remains, however, unclear whether random connectivity alone can account for specific olfactory percepts and actions, reproducible between individuals^30, 53^. Furthermore, identifying structure in the connectivity of the olfactory system has been impeded by the scarcity of single cell connectivity data available. While several studies addressed this challenge in insects^33, 52, 54–59^, comparable understanding of the statistics of ensembles of single olfactory bulb or piriform cortex output neuron projections in mammals is lacking^60^, despite recent advances in optical imaging-based single neuron tracing^61, 62^. To date, owing to the extensive intertwining of neural processes, axonal tracing studies reconstructed only dozens of individual bulb output neuron projections^13–16^, a number insufficient for analyzing the statistics of their branching patterns across multiple target regions. Resolving the question of structure in the olfactory system connectivity requires interrogating the projections of multiple brain regions at cellular resolution with high-throughput.

Here, we have mapped the projections of thousands of neurons originating from both the olfactory bulb and piriform cortex respectively using two high-throughput single-neuron projection mapping techniques, Multiplexed Analysis of Projections by sequencing (MAPseq) and Barcoded Anatomy Resolved by sequencing (BARseq)^34–38^. The high yield of these methods and the resulting sample sizes – two to three orders of magnitude more neurons probed than in previous studies – reveal the wiring of the olfactory system at unparalleled scale and resolution. Specifically, we asked whether a systematic relationship exists between the inputs from the olfactory bulb to the piriform cortex and the piriform cortex outputs. We identified matching input-output projection gradients along the A-P axis of the piriform cortex: piriform cortex outputs to specific brain regions match the collateral projections of their dominant olfactory bulb inputs to those same targets, thus supporting models of information processing that rely on structure in connectivity coordinated across brain regions.

### Mapping the brain-wide projections of olfactory bulb outputs using neuronal barcoding

To interrogate the organization of the olfactory bulb outputs, we used two single-neuron projection mapping techniques, MAPseq and BARseq^34–38^ (**Fig. 1a-c**; **Extended Data Fig. 1**; see Methods). MAPseq rapidly maps out the projections of thousands of neurons in a single animal^34, 35^ by uniquely labeling individual neurons in the source region (e.g. mitral cells in the bulb, **Fig. 1b**) using random RNA barcodes delivered via a Sindbis viral library. The rate of infection is kept low to ensure that most neurons are labeled by only one barcode^34, 35, 37^. These barcodes are then replicated and transported to the axonal branches. Projections to multiple brain regions can be determined by sequencing and matching axonal barcodes from these target regions (**Fig. 1c**) to the barcodes in the injection area (i.e. olfactory bulb).

**Figure 1.**
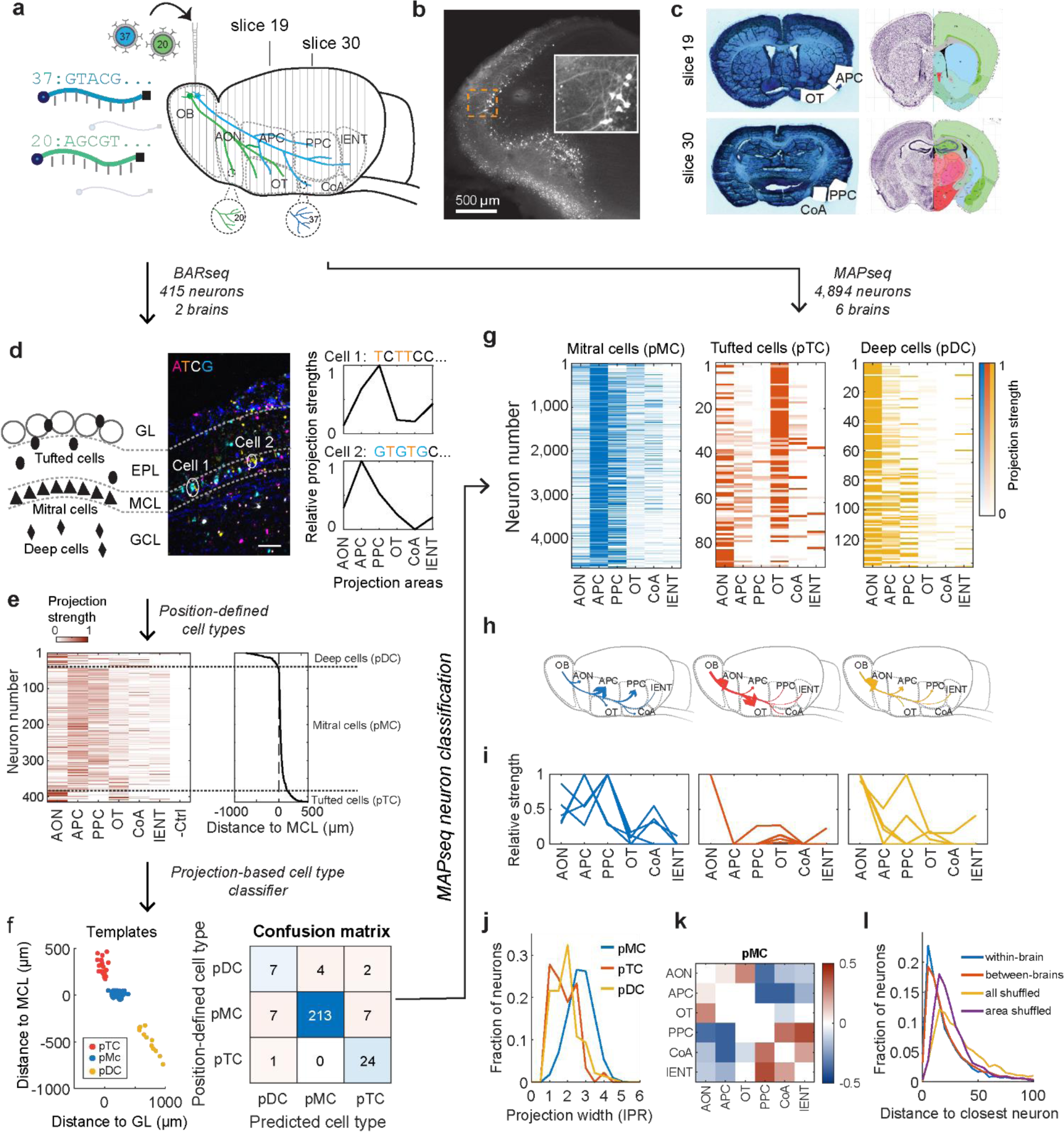
MAPseq and BARseq projection mapping of individual olfactory bulb neurons. (**a**) Schematics of the MAPseq strategy which uses RNA barcodes to label neurons and map their brain-wide projections. (**b**) Infection of mitral and tufted cells by Sindbis virus carrying the barcodes and a fluorophore (GFP). (**c**) Laser Capture Micro-Dissection of target brain regions from Nissl stained coronal sections and corresponding sections registered to the Allen Brain reference atlas. (**d**) Illustration of laminar positions of mitral, tufted, and deep cells (Left) and an example BARseq sequencing image of the barcoded cells (Center). The first several bases of the barcode sequences in two example neurons analyzed via BARseq and their projection patterns across 6 bulb target brain regions (Right). Scale bar = 100 µm. (**e**) Projection patterns of neurons (415 neurons, 2 mice) identified via BARseq and their soma locations relative to the mitral cell layer (MCL). Columns represent olfactory bulb projection target regions and rows indicate individual neurons. Cell identities based on soma positions are shown on the right. Projection strength of each barcoded neuron has been normalized so that the maximum strength is 1 in each neuron (row). (**f**) (Left) Soma positions of template neurons shown relative to MCL (y-axis) and to glomerular layer (x-axis) that were used to train the projection-based classifier. Neurons are colored by their identities based on laminar positions (tufted, mitral and deep cells). (Right) The classification confusion matrix of all three classes of neurons using the BARseq-based classifier versus the position-defined classes. (**g**)-(**i**) The projection patterns of all MAPseq analyzed neurons (**g**), their mean projection patterns (**h**), and five example neurons (**i**) of the three classes of bulb projection neurons identified via a BARseq-based classifier. In (**g**), columns represent projection brain regions and rows indicate individual barcoded neurons. Barcoded neurons are sorted by probability of cell type classification based on running their projection patterns through the classifier. (**j**) Distribution of the broadness of projections, as measured by Inverse Participation Ratio (IPR, x-axis) at brain region-level. (**k**) Pearson correlation between putative mitral cell (pMC) projections to different target regions. Only correlations that passed statistical significance after Bonferroni correction are shown. (**l**) Distribution of the city block distance between the projection patterns of each pMC identified using the BARseq-based classifier and the most similarly projecting pMC within the same brain (blue), across different brains (red), across all brains (6) sampled after shuffling all elements in the projection matrix (yellow), or after shuffling the neuron identities for each area separately (purple).

We collected tissue from the major bulb projection targets at 200 µm resolution along the A-P axis of the brain, including the anterior and posterior piriform cortex (APC/PPC), the anterior olfactory nucleus (AON), the olfactory tubercle (OT, olfactory striatum), cortical amygdala (CoA), and the lateral entorhinal cortex (lENT), which each play distinct functional roles^20, 63–69^. We carefully excluded fibers of passage from the lateral olfactory tract (**Supplementary Note 1**; Methods). Mitral and tufted somata are positioned across different layers of the bulb (mitral cell versus external plexiform layers) and further differ in size, morphology, intrinsic excitability, local wiring, and projection patterns^13, 14, 43, 70–77^. To distinguish mitral from tufted cells and possibly other bulb output neurons, we applied BARseq to a subset of the sampled brains. BARseq builds upon MAPseq, but additionally uses *in situ* sequencing to spatially resolve barcoded somata, thus enabling one to distinguish output neuron types barcoded in the injection area based on their laminar locations (**Fig. 1d**). Both BARseq and MAPseq have been extensively validated in other brain regions, and results have confirmed and extended those obtained by classical neuroanatomical methods, including retrograde labeling, bulk anterograde tracing, and single-cell tracing^34–37^. Previous reports also addressed other potential technical concerns, such as double labeling of neurons, distinguishing fibers of passage from axonal termini, and uniformity of barcode transport^34–37^. Our approach enabled mapping the projections of 5,309 bulb output neurons (415 neurons from two brains using BARseq, and 4,894 neurons from six brains using MAPseq), amounting to two orders of magnitude more individual neuronal projections than all previous single-cell studies of bulb connectivity combined.

### The distribution of olfactory bulb projections to their targets displays structured correlations

To determine the statistical structure of olfactory bulb projections, we first used BARseq to identify mitral and tufted cells based on the location of their somata (**Fig. 1d,e**). We defined putative mitral cells (pMC) as those with somata in the mitral cell layer and putative tufted cells (pTC) as those with somata in the external plexiform layer (see **Supplementary Note 2** for further discussion). Consistent with previous reports^8, 9, 13, 14, 47^, pTC predominantly projected to AON and/or OT, whereas pMC sent their axons broadly throughout the olfactory stream and displayed strongest projections to the anterior and posterior piriform cortex (APC and PPC, **Fig. 1e**). Interestingly, we also found a small, but distinct, third group of deep cells (pDC) in the granule cell layer that resembled deep short axon cells previously reported to innervate higher olfactory areas^78, 79^. Therefore, BARseq identifies and recapitulates known differences in the projection of olfactory bulb output neurons.

Although BARseq enables distinguishing neuronal types based on their somata positions, MAPseq can map the projections of a larger number of neurons. However, because the precise locations of the barcoded neurons are unknown in MAPseq, we cannot directly determine their identities. To take advantage of the higher throughput of MAPseq and distinguish across different neuronal types, we next used the BARseq dataset as template to classify 4,894 bulb output neurons whose projections we mapped using MAPseq. We trained and cross-validated a neural network-based probabilistic classifier to assign each neuron to the most likely type (mitral, tufted or deep cell) given its projections across the bulb target regions sampled. As ground truth for the classifier, we used the cell types defined by their somatic locations measured via BARseq (**Fig. 1f**; **Extended Data Fig. 2a-g**; **Supplementary Note 2**). In both BARseq and MAPseq experiments, viral injections were targeted to the mitral cell layer, allowing far more pMC to be infected than pTC and reflecting differences in cell density (**Fig. 1d**, **e**, **g**; see **Supplementary Note 2**). Indeed, running the classifier on neurons mapped using MAPseq, we identified 4,665 pMC, 90 pTC, and 139 pDC (**Fig. 1g-i**). The projection patterns of the classified neurons were distinct across types and were consistent with BARseq-identified neurons of the same type (**Fig. 1e**). Because of the prevalence of labeling in the mitral cell layer, we focused most of the subsequent analysis on a subset of pMC neurons (4,388) that was classified with high confidence (>85% classification accuracy; **Extended Data Fig. 2h**; **Supplementary Note 2**; Methods).

**Figure 2.**
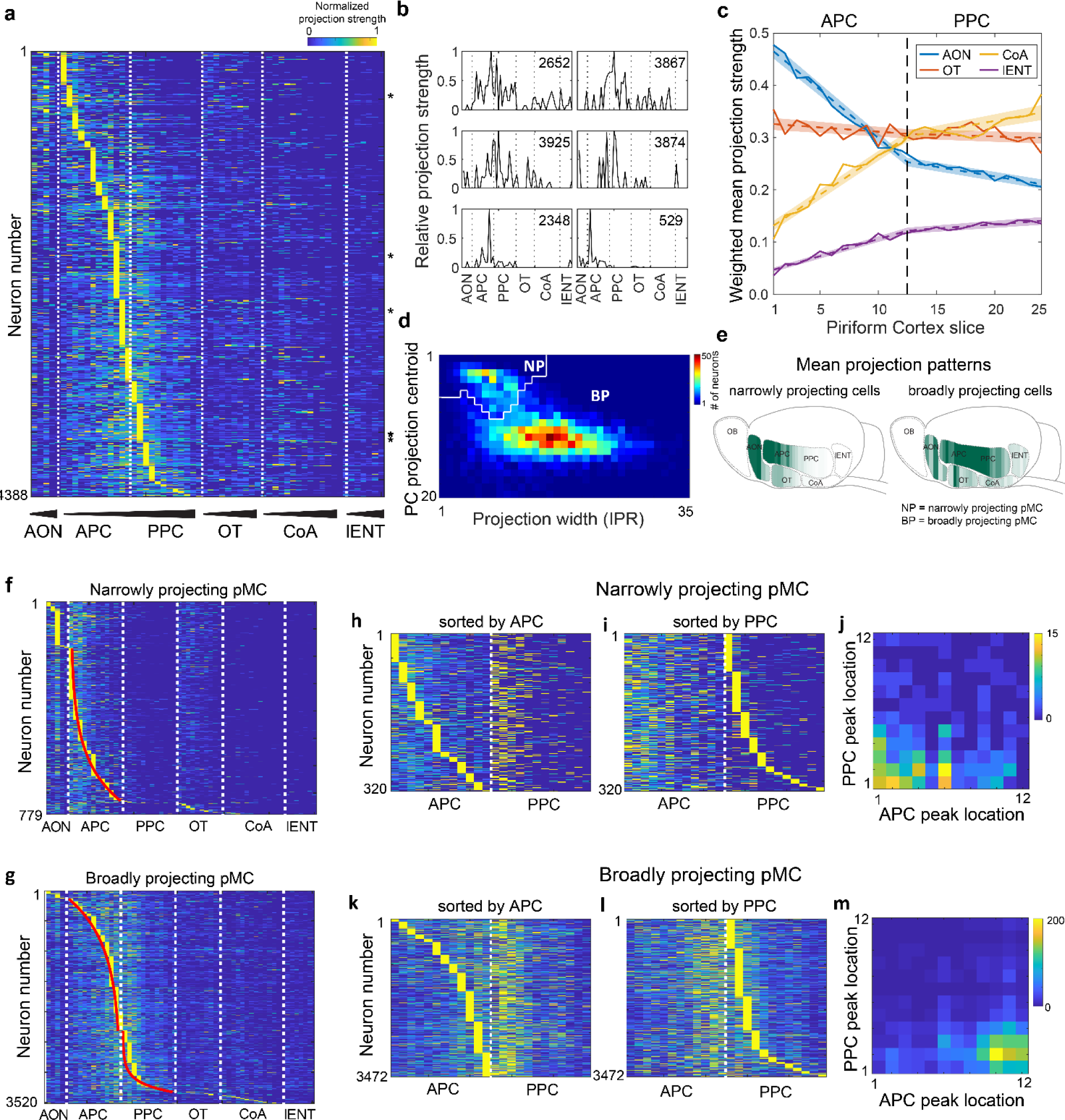
Spatial pMC projection gradients tile differentially the A-P axis of piriform cortex and are predictive of pMC co-projections to extra-piriform bulb target brain regions. (**a**) Projections patterns of pMC neurons at 200 µm resolution along the A-P of piriform cortex. Rows represent neurons and each column represents a single 200 µm section in the brain region indicated at the bottom. Within each region, slices are arranged by A-P positions, with the most anterior sections on the left. **(b)** Projection patterns of example pMC neurons. Projections are normalized by maximum barcode count for each neuron. Neuron numbers are indicated on top of each plot and correspond to the fiduciary marks shown in **(a)**. **(c)** Weighted mean projection strength for olfactory bulb neurons to four major extra-piriform targets as a function of location of piriform cortex co-innervation [the conditional probability of co-innervation P(target|PC location), solid lines]. Dashed lines/shaded areas show piecewise linear fits in APC and PPC with the 95% confidence interval obtained by bootstrap. (**d**) The distribution of neurons as a function of projection centroid and projection width (Inverse Participation Ratio, IPR). Two types of neurons form distinct clusters, the narrowly and broadly projecting pMCs (NP and BP). Watershed clustering identifies a separatrix (white line) for the distributions of these two cell populations. (**e**) Heat maps of mean projection patterns of narrowly and broadly projecting neurons. (**f**)(**g**) Projection patterns of narrowly (**f**) and broadly projecting (**g**) pMCs across target areas at 200 µm resolution. Neurons are sorted by peak projection positions across all slices. Red curves in APC show fits with exponential [Eq. (1)] and inverted exponential [Eq. (2)] functions for NP and BP cell projection strength maxima respectively along the A-P axis. Similarly, the red curve in PPC (**g**) shows an exponential fit for the BP cell projection strength maxima along the A-P axis. (**h-m**) For both narrowly and broadly projecting neurons, the locations of maxima of projections along the A-P axis within APC and PPC respectively are not correlated. (**h**,**i**) Narrowly projecting neurons sorted by the strength of their projection maxima along A-P axis of APC (**h**) and PPC (**i**). (**k**,**l**) Same for broadly projecting neurons. (**j**,**m**) Density plots of the distribution of peak projection positions of the same neurons within APC (x-axis) and their peak projection positions in PPC (y-axis). Colors indicate number of neurons.

Unlike pTC and pDC, individual pMC branched and projected broadly (**Fig. 1j**; inverse participation ratio, IPR of pMC is 2.71 ± 0.69 mean ± s.d., compared to 1.79 ± 0.66 for pTC and 1.90 ± 0.66 for pDC; p < 10^-10^ comparing pMC to pTC, as well as pMC to pDC, rank sum test and after Bonferroni correction). This raises the question whether the axons of individual putative mitral cells project randomly, or if instead the projections are structured in some way. For example, one group of mitral cells might project preferentially to one subset of targets, whereas another group might favor a different subset of targets. To test this hypothesis, we calculated the Pearson correlation between pMC projections to different bulb targets. Indeed, bulb projections to AON were correlated with bulb inputs to OT (**Fig. 1k** for pMC; r = 0.26, p < 10^-10^ after Bonferroni correction; **Extended Data Fig. 2i** for pTC and pDC), whereas projections to PPC were correlated with projections to the CoA and lENT (**Fig. 1k**; r = 0.31, p < 10^-10^ between PPC and CoA, and r = 0.39, p <10^-10^ between PPC and lENT after Bonferroni correction; **Extended Data Fig. 2i** for pTC and pDC). Consistent with the presence of correlations in the projections of individual pMC to bulb target regions, shuffling projections across pMC to each bulb target independently of other regions, significantly increased the minimal distance in projection space between pairs of pMC (**Fig. 1l**; p < 10^-10^, rank sum test). In agreement with previous work^17, 47^, we also identified differences in the connectivity of pMC originating from the dorsal versus ventral bulb aspects (**Supplementary Note 3**; **Extended Data Fig. 3a-c**), but no biases along the A-P axis of the bulb (**Extended Data Fig. 3d-f**). In summary, individual pMCs form several axonal branches that target multiple brain structures, but this connectivity is structured and highly selective, with specific combinations of brain regions targeted in a correlated manner.

**Figure 3.**
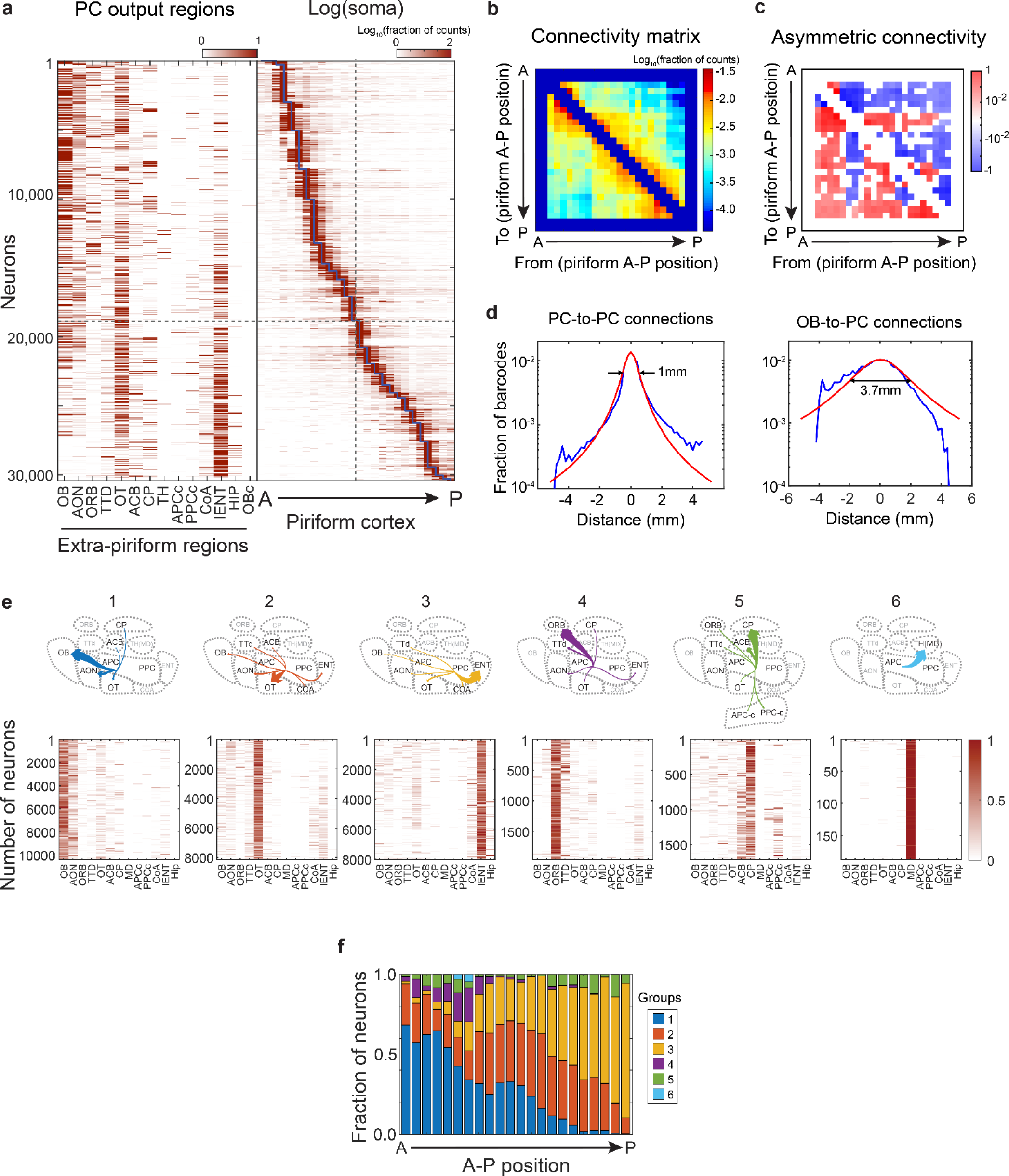
Projection patterns of piriform cortex output neurons are systematically organized along the A-P axis with respect to their extra-piriform targets. (**a**) (Left) Projection patterns of piriform cortex output neurons to extra-piriform brain regions (Supplementary Table 4) and (Right) within the piriform cortex along the A-P axis. Projection density is color-coded on a log scale. In the piriform cortex, for a given barcoded neuron, the A-P position with the most barcode counts is taken as the location of the soma. (**b**) Mean projection strengths (log scale) of projections from somata at the indicated locations (x-axis) to the specific A-P positions within the piriform cortex (y-axis). (**c**) Differences in reciprocal projections between two A-P positions in the piriform cortex obtained by calculating the difference between the connectivity matrix (b) and its transpose. Blue indicates stronger projection in the posterior direction, and red indicates stronger projection in the anterior direction. (**d**) (Left) The strength of intra-piriform projections relative to their soma locations (blue). Red line indicates fit using an inverse power law model (Methods). The density of projections decreases by half at about 0.5 mm from soma location (maximum density of barcodes), making the projections width equal to 1 mm at 50% density (arrows). Contribution from dendritic neuropil of barcoded neurons was minimized by removing slices adjacent to the peak of barcode molecule counts (Methods). (Right) the same distribution obtained for pMC is substantially broader. (**e**) Mean projection patterns (Top) and the projection patterns of individual neurons (Bottom) of groups of piriform cortex output neurons to extra-piriform target brain regions. (**f**) Fraction of neurons belonging to each group at the indicated A-P positions within the piriform cortex. The color codes used are the same as in (**e**).

### Olfactory bulb-to-piriform cortex connectivity along the A-P axis is predictive of the olfactory bulb co-innervation of extra-piriform target regions

Anterior and posterior piriform cortex each span ∼2.5 millimeters along the anterior-posterior (A-P) axis. Axons of individual pMC targeting the piriform cortex branch extensively and co-innervate several functionally distinct extra-piriform target regions. To better understand the logic of these projections, we investigated whether the finer spatial pattern of individual bulb neuron projections within the piriform cortex is predictive of their co-projections to extra-piriform targets. To test this hypothesis, we computed the conditional probability *P(target|PC location)* for each barcode sequence to be found in the four extra-piriform regions sampled (*target*), conditioned on the probability of finding the same barcode sequence at a given piriform cortex location along the A-P axis (at 200 µm resolution; **Fig. 2a-c**; **Extended Data Fig 4a-c**; **Supplementary Note 4**; Methods). Hence, this metric describes the rate (projection strength) at which bulb output neuron axons co-innervate the other four extra-piriform bulb major targets as a function of which location in the piriform cortex they innervate.

**Figure 4.**
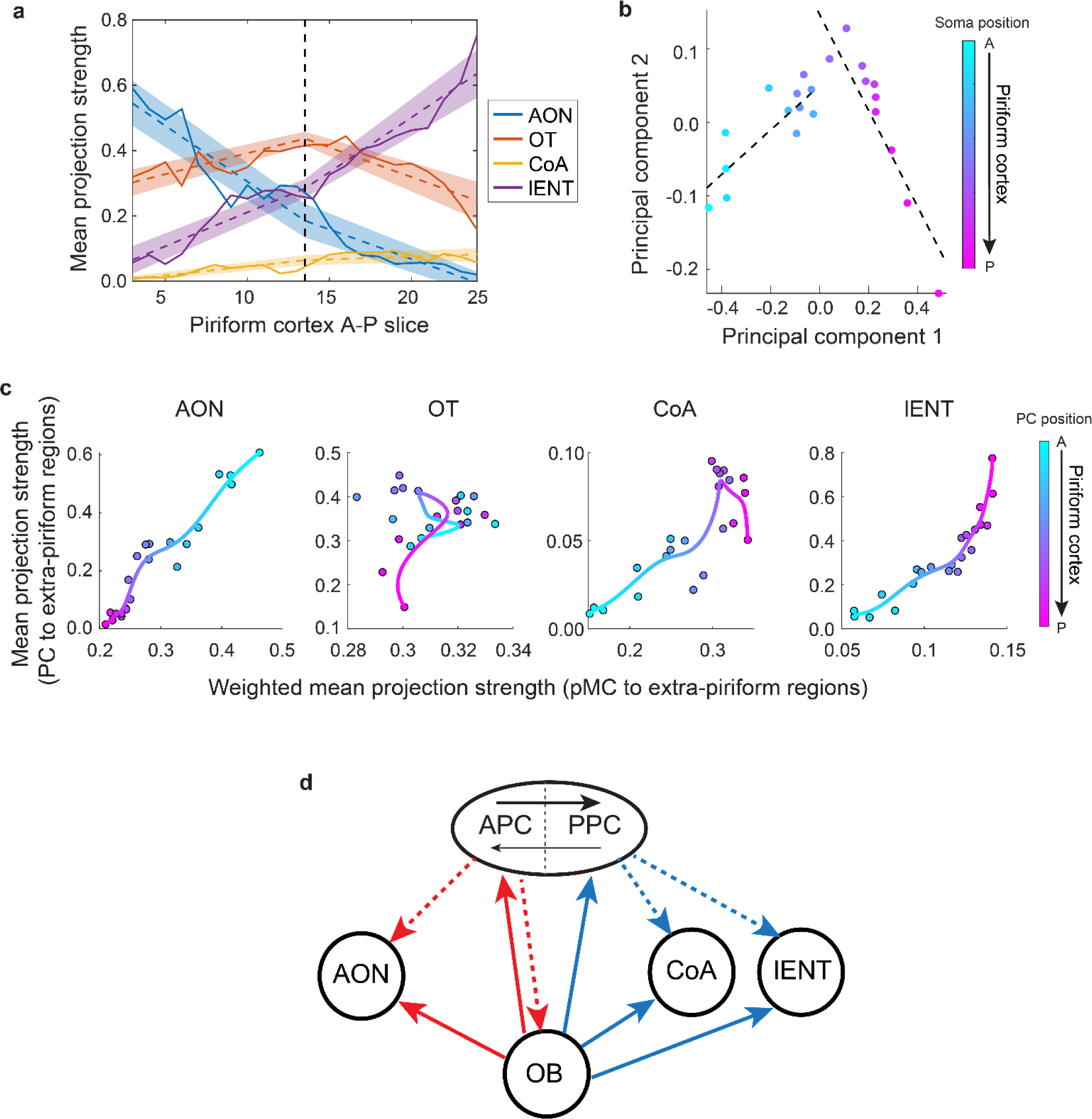
Matched input and output circuit motifs in the piriform cortex. (**a**) Mean projection patterns of piriform cortex output neurons at the indicated A-P position of barcoded somata in the piriform cortex. Dotted lines indicate linear fits and shaded areas the range of fits from bootstrap. (**b**) Mean loadings for the first two principal components of the mean projection strengths of piriform output neurons to AON, CoA, lENT and OT sampled at the indicated A-P positions in the piriform cortex. Dotted lines indicate linear fits for APC and PPC. (**c**) Mean projection strengths of piriform projection neurons to extra-piriform target regions, organized by the location of their somata along the A-P axis of the piriform cortex (y-axis) plotted against the mean projection strengths of pMC neurons to extra-piriform bulb target regions weighted by projections to a particular A-P position in piriform cortex, P(target|PC location) (x-axis). Colors indicate A-P positions in the piriform cortex. (**d**) Cartoon schematics of the parallel olfactory circuits engaging the olfactory bulb-to-piriform inputs, piriform cortex (APC and PPC) outputs and extra-piriform bulb target regions (AON, CoA and lENT) sampled in this study.

Remarkably, *P(target|PC location)* displayed orderly, close to linear, variation as a function of the piriform cortex location for AON, CoA and lENT, but not OT. For example, pMC projections to the AON were stronger in neurons that also innervated more strongly the anterior portion of the piriform cortex than those projecting more strongly to the posterior portion (**Fig. 2c**, ρAON = -0.99, p = 1.9 x 10^-23^, Spearman correlation after Bonferroni correction; **Extended Data Fig. 4b,c**). Conversely, bulb inputs to CoA and lENT were correlated with strong projections to the posterior portion of the piriform cortex (**Fig. 2c**, for CoA ρ_CoA_ = 0.95, p = 6.2 x 10^-13^; for 1ENT ρ_1ENT_ = 0.98, p = 2.7 x 10^-17^, Spearman correlation after Bonferroni correction; **Extended Data Fig. 4b,c**). The slopes of *P(target|PC location)* along the A-P axis were characteristically distinct for different brain regions (AON vs. lENT vs. CoA, **Fig. 2c**) and, furthermore, were different across APC and PPC (**Fig. 2c**, p < 0.005 for AON, CoA and lENT; p-values calculated using bootstrap and Bonferroni correction). The observed inter-dependence of projections has not previously been reported, presumably because previous datasets were too small for these patterns to become apparent (**Extended Data Fig. 4d**).

One possible explanation for these relationships is that projections simply “disperse” locally around a target area^3, 80–82^. This model predicts that pMC projections to OT, which neighbors APC, but not PPC, would also show a gradient along the A-P axis of the piriform cortex. However, such a gradient was not observed, suggesting that the locality model is not sufficient to account for our findings (**Supplementary Note 4**). These relationships were robust across individuals and were corroborated at the population level using CAV-2 Cre retrograde labeling (**Extended Data Fig. 5a, b**; **Supplementary Note 5**). Thus, the distribution of olfactory bulb co-innervation of extra-piriform regions varies in a systematic and specific manner with the strength in projection along the A-P axis of the piriform cortex.

### Projections of narrowly and broadly projecting pMCs differentially tile the A-P axis of the piriform cortex and innervate distinct domains of extra-piriform bulb target regions

Further analysis identified two distinct pMC populations — “narrowly projecting (NP)” and “broadly projecting (BP)” — according to the width and location of projections to both the piriform cortex and extra-piriform brain regions (**Fig. 2d-g**; **Extended Data Fig. 6a**, see **Supplementary Note 6** for discussion on their potential identity). The narrowly projecting population was smaller (∼20%) and displayed a more compact innervation pattern (IPR = 7.8 ± 2.8, mean ± std; Methods), whereas the remaining (∼80%) broadly projecting pMC formed more diffuse projections (IPR = 15.2 ± 5.6, mean ± std, **Fig. 2d**). Narrowly projecting cells targeted more strongly the anterior part of anterior piriform cortex (**Fig. 2e, f**), whereas the broadly projecting population innervated predominantly the caudal part of anterior piriform cortex and the boundary between the anterior and posterior piriform cortex (**Fig. 2e, g**). Both narrowly and broadly projecting populations tiled the A-P axis of anterior piriform cortex, but their maximum projection density locations followed different distributions (**Fig. 2f, g**). We observed a similar coverage without gaps of the posterior piriform cortex A-P axis by broadly projecting, but not narrowly-projecting cells projection maxima (**Fig. 2f, g**). When sorting the narrowly projecting cells according to the location of their peak projection in the anterior piriform cortex, these cells formed a distribution well-described by an exponential function (red curve in **Fig. 2f**; Methods):

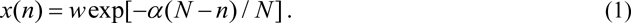

where *x* is the distance of the projection maximum from the APC anterior boundary, *w* the width of anterior piriform cortex extent along the A-P axis, *n* the cell’s rank according to the sorted location, *N* the total number of cells in the narrowly projecting population, and *α* = 3.8 the fitting parameter. The projection density of narrowly projecting cells as a function of A-P location in the anterior piriform cortex can be represented by: *ρ*(*x*) =| *dn* / *dx* |= *N* / *α x*. Thus, the density of inputs from narrowly projecting cells diverges near the anterior APC boundary (i.e. when *x* → 0, density of inputs is very high). Interestingly, an example of divergence in input sampling by a sensory system is provided by the logarithmic magnification factor found in the primary visual cortex with respect to the representation of retina^5, 83^. However, in the visual system, cortical areas similar in size take inputs from retinal regions shrinking towards fovea, while in the olfactory system, similar numbers of narrowly projecting cells target increasingly smaller regions of the anterior piriform cortex. Thus, in terms of sampling strategy, the olfactory cortex representations appear to be inverted compared to retinotopy.

The tiling of the anterior piriform cortex by broadly projecting cells was best described by an inverted exponential relationship given by (red curve in **Fig. 2g**):

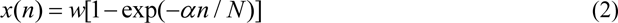

with *α* = 2.9. Equation (2) shows that the density of piriform cortex innervation for broadly projecting cells increases toward the posterior APC boundary, i.e. when *x* → *w*, *ρ*(*x*) = *N* / *α* (*w* − *x*) (**Fig. 2g**).

Finally, the tiling of the posterior piriform cortex by broadly projecting cells was also accurately described by an exponential distribution, similar to the coverage of the anterior piriform cortex by the narrowly projecting cells [Eq. **Error! Reference source not found.** with *α* = 6.5].

For both broadly and narrowly projecting cells, the locations of maximum projections of the same neuron along the A-P axis in the posterior piriform cortex were not correlated with the locations of maxima in the anterior piriform cortex (**Fig. 2h-m**, **Supplementary Note 6**), raising the possibility that local neural circuits in these two piriform cortex subdivisions perform different computations and extract distinct features of odor inputs. Furthermore, the distributions of projection peaks along the A-P axis in each of the extra-piriform bulb targets sampled were distinct for the narrowly and broadly projecting populations (**Extended Data Fig. 6b**): broadly projecting cells projected more posterior in OT (p < 10^-10^), CoA (p < 10^-10^), and lENT (p <10^-10^), but more anterior in AON compared to narrowly projecting cells (p < 10^-10^).

We investigated whether differences in projection between the narrowly and broadly projecting cells could account for the gradients described along the A-P axis of the piriform cortex. Area level correlations between projections to the piriform cortex and extra-piriform targets were weaker when analyzing narrowly and broadly projecting neurons independently versus all pMC jointly (**Fig. 1k**, **Extended Data Fig. 6c**), suggesting that these differences partially accounted for the brain region-level correlations observed. Nonetheless, at single slice (200 µm) resolution, the strength of co-projection to AON was higher for both narrowly and broadly projecting neurons targeting the anterior vs. posterior part of the piriform cortex. Similarly, the strength of co-projection to CoA and lENT was higher for narrowly and broadly projecting cells targeting the posterior vs. anterior part of the piriform cortex, forming orderly gradients (**Extended Data Fig. 6d, Supplementary Table 3**). Thus, the relationships described above (**Fig. 2c**) between the A-P spatial patterns of individual bulb neuron projections within the piriform cortex and their extra-piriform co-projections were preserved for both populations of pMC.

In summary, the olfactory bulb-to-piriform cortex connectivity along the A-P axis is predictive of the bulb co-innervation of extra-piriform target regions across different cell types (mitral vs. tufted cells^13, 14^), between two populations of pMC (narrowly and broadly projecting), and within the narrowly and broadly-projecting populations. Thus, these systematic relationships between projections to the piriform cortex and extra-piriform areas likely reflect a general wiring principle in the olfactory system.

### Piriform cortex output neurons distributed along the A-P axis project to different target regions

The correlations identified between the olfactory bulb projections along the A-P axis of the piriform cortex and the strength of their co-innervation of extra-piriform targets, together with reports on asymmetric rostro-caudal intra-piriform inhibition^84, 85^ and biases in the piriform cortex projections^86^, raised the possibility that downstream piriform cortical circuitry is also organized along A-P spatial gradients. To test this hypothesis, we used MAPseq to analyze the projections of the piriform cortex output neurons spanning 5 mm along the A-P axis (30,433 neurons from five mice, **Fig. 3a**). We sampled the piriform cortex (APC and PPC, **Fig. 3a**, *Right*) and 14 of its projection target regions (**Supplementary Table 4**, **Fig. 3a**, *Left*), identifying the locations of barcoded somata of the piriform cortex outputs (blue line in **Fig. 3a**, *Right*), their intra-piriform projections at 200 µm resolution and their projections to extra-piriform targets (Methods). Several features of intra-piriform connectivity (high connectivity within APC and PPC, **Extended Data Fig. 7a**, APC-to-PPC projection bias, **Fig. 3b, c**) and brain-wide organization of piriform cortex output neurons (**Extended Data Fig. 7b-e**) observed using MAPseq matched previous reports, confirming the robustness of our approach^49, 87^ (**Supplementary Note 7**).

Further analyses revealed novel intra-piriform local connectivity and piriform cortex output projection motifs. Specifically, we find that intra-piriform connections, while fairly long-range, decreased with distance, following an inverse power law (**Fig. 3d**). The density of connections dropped by approximately an order of magnitude at distances from somata of about 2 mm. Overall, the decay in intra-piriform projection strength was more substantial for the anterior than for posterior oriented connections in agreement with a previously observed anterior-to-posterior projection bias^49, 87^ (**Fig. 3a-c**). The fact that mitral cell barcodes were successfully transported along several millimeters from the olfactory bulb (**Fig. 2a**, **3d**), as well as previous demonstrations of long-distance (> 10 mm) transport of barcodes from other brain structures^34, 36, 37^, argued against the possibility that the observed locality of intra-piriform connectivity was an artifact of barcode transport. Thus, a distinctive feature of intra-piriform connectivity is its power-law decay as a function of distance from somata. While this contrasts the sharper exponential decay observed in other brain regions^82, 88, 89^, it points to a local organization of the piriform cortical circuit.

We further investigated whether the piriform cortex outputs are systematically organized along its A-P axis with respect to their extra-piriform targets. We separated the piriform cortex output neurons into six groups based on their extra-piriform projection targets using Louvain community detection (**Fig. 3e**; **Supplementary Note 8**; Methods). These groups were indeed differentially distributed with respect to the A-P position of their somata, had distinct intra-piriform connectivity and innervated specific combinations of brain regions, each having a dominant target region (**Fig. 3f**; **Extended Data Fig. 8a**-**f**; **Supplementary Note 8**). For example, groups 1 and 4 projected most strongly to the olfactory bulb and orbitofrontal cortex (ORB), respectively, and were enriched in the anterior piriform cortex. In contrast, neurons of group 3 projected most strongly to the lENT and were predominantly found in the posterior piriform cortex (**Fig. 3f**). These differences in soma distribution for the six groups were consistent across animals (**Extended Data Fig. 8c**). In summary, piriform cortex output neurons that project to different sets of target regions are enriched at specific locations along the A-P axis.

### Matching projection gradients of piriform cortex inputs and outputs along the A-P axis

We investigated whether the piriform cortex outputs form projection gradients reminiscent of the olfactory bulb-to-piriform cortex inputs. Consistent with the overall organization of the olfactory bulb-to-piriform cortex projections, the strength of piriform cortex output projections to AON, lENT and CoA varied along the A-P axis of the piriform cortex (**Fig. 4a**). The mean strength of piriform projection to AON decreased systematically from the anterior to the posterior end of the piriform cortex (Spearman correlation ρ = - 0.96, p = 2.0 ×10^-12^ after Bonferroni correction), while the mean strength of the piriform cortex projection to lENT and CoA increased (Spearman correlation ρ = 0.77, p = 7.5 × 10^-5^ for CoA and ρ = 0.97, p = 1.2 × 10^-14^ for lENT after Bonferroni correction). In contrast, piriform cortex projections to OT appeared enriched near the APC / PPC boundary but lacked an overall trend across the whole PC (Spearman correlation ρ = -0.11, p = 1 after Bonferroni correction). The observed piriform cortex gradients were further supported by retrograde bulk labeling experiments using fluorescent microbeads (RetroBeads) (**Extended Data Fig. 9a-c**; Spearman correlation between the A-P positions of neurons and projection strengths were ρ = -0.94, p = 2.0 × 10^-5^ for AON, ρ = 0.92, p = 1.7 × 10^-5^ for CoA, and ρ = 0.74, p = 4.4 × 10^-4^ for lENT after Bonferroni correction; n = 2 animals for AON and CoA and n = 3 animals for lENT). In addition, performing principal components analysis (PCA) on the projection strengths of the piriform cortex output neurons across these four extra-piriform areas revealed a sharp change in slope at the APC-PPC boundary (**Fig. 4b**, Methods). Thus, piriform cortex outputs to different brain regions form monotonic projection gradients along the A-P axis which follow different slopes in the anterior and posterior piriform cortex.

Since both the olfactory bulb-to-piriform cortex inputs and the piriform cortex output projections appear spatially ordered along the A-P axis of the piriform cortex (**Figs. 2c**, **3f**), we next asked whether they match. Strikingly, along the A-P axis, the mean strengths of piriform cortex projections to AON, CoA, and lENT strongly correlated with *P*(*target|PC location*) of pMC projections to each of these brain regions (**Fig. 4c**, Spearman correlation ρ = 0.95, p = 1.6 × 10^-5^ for AON, ρ = 0.73, p = 6.7 × 10^-4^ for CoA, and ρ = 0.97, p = 1.4 × 10^-5^ for lENT after Bonferroni correction). For example, neurons in the anterior part of piriform cortex (**Fig. 4c**, left panel, cyan dots) project strongly to the AON (y-axis), and the pMC neurons that target this part of piriform cortex also target strongly the AON (x-axis). Conversely, neurons in the posterior part of the piriform cortex (magenta dots) project weakly to the AON, matching the weak AON projections from pMC neurons that also target the posterior part of the piriform cortex. Similar matching relationships between the strength of piriform cortex output projections and of co-innervation by pMC inputs were also seen for CoA and lENT, but with reversed mapping along the A-P axis (i.e. strong projections from and to the posterior part of piriform cortex respectively). No robust relationship was observed for projections to the OT when performing the same analysis (Spearman correlation ρ = 0.03, p = 1 after Bonferroni correction).

Therefore, a given piriform cortex output neuron appears to be wired up such that the strength of its projection to AON, lENT or CoA matches the strength of its dominant bulb input’s co-projection to the same target brain region (**Fig. 4d**). These matching gradients in the inputs and outputs of the piriform cortex, consequently, may connect co-innervation targets of individual olfactory bulb cells (i.e. piriform and extra-piriform areas), thus completing parallel circuits which involve direct and indirect pathways.

## Conclusions

We investigated the wiring logic of the olfactory system using barcode sequencing-based mapping of thousands of individual olfactory bulb and piriform cortex output neurons. Our mapping strategy confirmed previous reports on the organization of bulb projections (i.e. mitral vs. tufted cells)^8,9, 13, 14, 47, 90^, as well as biases in intra-piriform connectivity^49, 87^ and piriform cortex output projections^86, 91, 92^ (**Figs. 1**, **3**). In addition, we identified novel and robust olfactory bulb projection gradients (**Fig. 2**), as well as groups of piriform cortex output neurons (**Figs. 3, 4**), which differentially tile the anterior-posterior axis of the piriform cortex and co-innervate specific sets of brain regions. Taken together, these findings indicate the presence of spatially structured connectivity in the mouse olfactory system, both at the level of non-random combinations of olfactory bulb and piriform cortex target regions and, at finer resolution, within the piriform cortex. Furthermore, the projection gradients we identified support matched, parallel input-output piriform cortex circuit motifs which co-innervate select extra-piriform bulb targets (**Fig. 4**) and enable direct and indirect processing pathways.

Previous studies have concluded that the olfactory bulb sends broad and, potentially, random projections to the piriform cortex^8, 9, 11, 13, 15, 16, 47^. These observations have implications for the nature of computations performed by the olfactory circuit. Our findings do not necessarily contradict a framework of broad connectivity within the piriform cortex. Indeed, broad projections may contain spatial biases, if projection widths are comparable or less than the spatial extent of the target area (i.e. piriform cortex). Rather, our data are consistent with a model in which the olfactory bulb-to-piriform cortex connectivity contains both distributed and spatially organized components. Our data do, however, rule out the hypothesis that the olfactory bulb-to-piriform cortex connectivity is random and contains no spatial structure. Indeed, the axons of bulb output neurons form orderly representations in the piriform cortex, as defined by distinct and reproducible patterns across different cell types (mitral vs. tufted cells) and subpopulations (narrowly and broadly projecting cells) along the A-P axis and by their co-projections to other brain regions. These findings, in conjunction with previously observed gradients in the piriform cortex inhibitory connections^84, 85^, point to a model in which the A-P location within the piriform cortex may reflect a feature or a set of features of olfactory bulb responses that are important for cortical processing. These features appear to *not* be represented topographically in the bulb. Furthermore, as it follows from the uncorrelated nature of narrowly and broadly-projecting cells tiling in the anterior and posterior piriform cortex (**Fig. 2j, m**), the features represented may differ in these cortical subdivisions. Within this framework, the olfactory bulb-to-piriform cortex projections are organized topographically while subject to developmental variability. Therefore, in this model, the olfactory cortex is topographically organized, similarly to other sensory cortical areas (i.e. visual, auditory, somatosensory cortex) according to stimulus features that need further investigation.

The spatial organization of connectivity we report here (**Fig. 2c**) could, in principle, be understood based on spatial locality in connectivity^3, 80–82^: axons tend to form connections to nearby neurons/areas and, consequently, the density of connections decreases as function of distance. This may emerge, for example, because brain regions are arranged so as to minimize the total length of connections^3, 93^. As such, the locality rule is the default model to apply to connectivity. We find however that several connectivity features cannot be explained by this purely geometric model. First, the connectivity to the olfactory tubercle appears to be rich and flat, despite distance to the tubercle varying substantially along the piriform cortex A-P axis. Second, the slope of spatial dependence of connection probability differs across olfactory bulb target regions (lENT vs. CoA vs. AON; **Fig. 2c**). Third, at the boundary between the anterior and posterior piriform cortex, we observe sudden changes in slope (**Fig. 4b**) for projections to different extra-piriform olfactory bulb target regions. Finally, the choice of dominant projection of the piriform cortex output neurons cannot be explained by proximity of the target region to the soma location within the piriform cortex (e.g. projections to the bulb are stronger than those to AON; **Fig. 3e**). Overall, these features may be better explained instead by a function-based topographical model. Of note, the function- and location-based connectivity models are not mutually exclusive. Indeed, since circuit function is expected to define connectivity, the geometry and locations of connected regions may be determined based on connection length minimization^3, 93^. As a consequence, location-based connectivity features may be explained by the functional needs of the circuit. While the geometric and functional connectivity models yield similar first order spatial biases in projection, the functional model more accurately explains details in the data, such as differences in the slopes of projection variations (**Fig. 2c, 4a**).

We also find that the density of connections within the piriform cortex decays as a power-law function of distance from somata (**Fig. 3d**). This is somewhat slower than the exponential decay observed in some brain regions^82, 88, 89^. However, intra-piriform connectivity does not extend with equal density across the entire piriform cortex. For example, the density of connections decays by more than an order of magnitude for loci separated by 2 mm along the A-P axis, indicative of local organization in intra-piriform connectivity. This finding, based on sequencing the projections of ∼30,000 piriform cortex neurons in five brains, contrasts previous suggestions derived from a small sample of neurons^24, 25^. The local nature of connectivity argues strongly for spatial organization of functionality within the piriform cortex. According to classical proposals^11, 20, 94^, one key role of the piriform cortex is to implement pattern completion by linking neurons responding to the same stimuli. If relevant parts of these responses are distributed across the piriform cortex, as suggested by the random projection model, the associative intra-piriform circuit is expected to be highly distributed and long-range. Instead, our observations are consistent with a model in which associative connectivity within piriform cortex is organized locally and integrates locally activated ensembles of neurons.

Further supporting the hypothesis of spatial structure in the piriform cortex circuit function, we found that inputs from the bulb correlate with specific piriform cortex output projections in the way they vary along the A-P axis. Indeed, using the A-P axis as reference, our experiments identified systematically matched input-output circuit motifs (**Fig. 4**). The monotonicity of the piriform cortex input and output projection gradients, their specificity with respect to different extra-piriform bulb target regions (i.e. AON vs. CoA or lENT), and the differences observed between cortical subdivisions (APC vs. PPC) are further indicative of spatially ordered olfactory representations. These results open venues for further understanding the molecular mechanisms underlying the emergence of these gradients during development. Interestingly, the circuit motifs identified here are reminiscent of Residual Neural Networks (ResNet)^95^ architectures. They mirror similar motifs reported in both sensory networks, such as the topographically aligned retino-tectal vs. retino-thalamo-cortico-tectal projections^96–98^ and in the limbic system (thalamo-amygdala vs. thalamo-cortico-amygdala projections^99, 100^ or entorhinal cortex-CA1 vs. entorhinal cortex-to-dentate gyrus-CA3-CA1^101, 102^). This architecture enables both implementing different computations on inputs from the sensory periphery (e.g. olfactory bulb, retina) via parallel streams, and cross-talk and comparisons across functional streams. An intriguing possibility is that these input-output circuit motifs process different stimulus variables (i.e. sensory vs. contextual or sensorimotor information)^20, 103^. Future work may relate the odorant receptor molecular identity of glomeruli to the brain-wide projections and responses of these inter-connected bulbar and piriform cortex outputs during behavior.

In summary, we find that the piriform cortex outputs to specific brain regions match the collateral projections of their dominant bulb inputs to those same targets, thus forming a series of input-output circuit motifs organized along the A-P axis of piriform cortex. Each motif includes parallel pathways resembling the ResNet network architecture, a connectivity motif often found across the brain. Thus, while the olfactory cortex map does not represent the olfactory bulb topography, piriform circuits may extract odor features that are processed in a spatially organized manner, akin to other sensory modalities. Our findings provide a foundation for the functional interrogation of olfactory processing that relies on the presence of structure in the bulb-to-piriform inputs and the piriform cortex connectivity.

## Methods

### Barcoded Sindbis viral library

A barcoded Sindbis virus library (JK100L2, Addgene plasmid #79785) was generated and used for MAPseq and BARseq experiments as described previously^34–37^. *In vitro* sequencing of the viral library revealed that the library contained at least 8 million different barcode sequences. No barcode sequence was found more than once in the 5,309 OB output neurons sampled via MAPseq (6 brains) and BARseq (2 brains). Only 17 barcodes were identified more than once in the 30,433 PC output neurons sampled via MAPseq (5 brains).

### Animals

3 C57BL/6J of 8-10-week old male mice (Jackson Laboratory) were used for MAPseq and BARseq experiments (8 for mapping the olfactory bulb output projections, and 5 for mapping the piriform cortex outputs). In addition, 11 mice were used for experiments cross-validating the projection patterns observed via MAPseq (5 for fluorescent microbeads injections and 6 for CAV2-Cre viral labeling, respectively). All animal procedures conformed to NIH guidelines and were approved by the Institutional Animal Care and Use Committee of Cold Spring Harbor Laboratory. A list of all animals used is provided in **Supplementary Table 1**.

### Viral labeling of olfactory bulb (OB) output neurons

Mice were anesthetized using oxygenated 4% isoflurane and maintained with oxygenated 0.7-1.0% isoflurane throughout the surgery. Mice received subcutaneous injection of meloxicam (2 mg/kg), dexmetasone (1 mg/kg) and baytril (10 mg/kg) immediately before and after surgery. Lack of pain reflexes was monitored throughout the procedure. Mice were positioned such that the skull dorsal surface is horizontal. Animals were mounted in a stereotaxic frame (David Kopf Instruments), and small craniotomies were performed into the skull above the olfactory bulb, such as to enable injections at multiple penetration sites along the anterior-posterior (A-P) axis. We injected 50 nl Sindbis virus (1:3 diluted) at two depths of 100 µm and 200-250 µm (from brain surface) at four penetration sites along the A-P axis of the dorsal aspect of the bulb (0.8 mm, 1.2 mm, 1.7 mm, 2.2 mm anterior to the blood vessel between OB and prefrontal cortex, 0.7-1.0 mm lateral from midline). In a subset of animals (4 mice), we also injected 70 nl (1:3 diluted) Sindbis virus across two depths at 3 penetration sites on ventral aspect of the bulb (0.75 mm anterior, 1.2-1.1 mm lateral, 1.2 and 1.45 mm deep; 1.3 mm anterior, 1.0-1.1 mm lateral, 1.0 and 1.3 mm deep; 1.8mm anterior, 0.8-0.9mm lateral, 1.0 and 1.3mm deep) (**Extended Data Fig. 1a**). For the BARseq experiments (2 mice), we injected 70 nl viruses at 200-250 µm from brain surface into six sites, spaced 500 µm apart along the A-P axis of the dorsal aspect of the bulb (**Extended Data Fig. 1a**). We slowly pressure-injected (Picospritzer II, Parker Hannifin) the virus within 10-15 mins via a glass pipet at each depth and waited for 5 mins afterwards. After injection, we applied Kwik-Cast (Word Precision Instrument) to cover the exposed craniotomy and sealed the skin above using tissue adhesive (3M vetbond). Mice were allowed to recover, while waiting for expression of the virus, and euthanized 44-48 hours post-injection.

### Cryo-sectioning and laser microdissection for MAPseq of OB outputs

The brain was immediately extracted from the skull, flash-frozen on dry ice and temporarily stored at -80°C before cryo-sectioning. The whole brain was sliced into 200 µm coronal sections along the A-P axis using a Leica CM 3050S cryostat set at -10°C. To avoid cross-contamination, we used a fresh unused part of a blade to cut each slice and cleaned the brush and the holding platform with 100% ethanol between slices. Each brain slice was melted on a steel frame PEN-membrane slide (Leica microsystems, Cat #11600289) pre-coated with Poly-L-Lysine (Sigma). OB slices cut from brains injected with the Sindbis virus on dorsal OB side only (brains YC61&65) were processed slightly differently. These were melted onto a clean microscope glass slide, rapidly frozen on dry ice and later hand-dissected using cold scalpel while keeping the section frozen. Each brain slice placed on a PEN-membrane slide was immediately processed using following procedure as previously described^37^: fixation in ice-cold 75% EtOH for 3 minutes, one rapid rinse in Milli-Q water (Millipore) at room temperature, staining in 0.25% Toluidine Blue O (Sigma-Aldrich, MO) in Milli-Q water for 30 seconds at room temperature, followed by three quick rinses in Milli-Q water and two post-fixation rinses in 75% EtOH (3 mins each) at room temperature. After processing OB slices, we changed to fresh buffers for the rest of brain slices containing the OB target regions so as to avoid contamination of the target regions by barcodes from the injection site. After fixation and staining, we placed each brain slice on a PEN-membrane slide to dry for 30-60 minutes in a vacuum desiccator, with desiccant at bottom, at room temperature before attaching a second fresh PEN-membrane slide on top of the brain slice. The two PEN-membrane slides, sandwiching the brain slice in the middle, were taped together to prevent the brain slice from falling off or curling up as it continued to dry in the vacuum desiccator overnight.

Dorsal and ventral aspects of OB and target brain regions was laser micro-dissected from each brain slice on a Leica LMD7000 using the Allen mouse Brain Atlas as reference. Each region of interest dissected from every brain slice was collected and transferred into a single homogenizing tube with a homogenizing bead, added with 100 µl lysis solution of RNAqueous-Micro total RNA Isolation kit (Thermo Fisher Scientific), immediately frozen on dry ice, and stored at -80°C as previously described^37^. To assess cross-sample contamination, we also collected, as negative controls, tissue from brain regions known to receive no direct input from the OB, such as contralateral piriform cortex, ipsilateral primary motor, somatosensory and visual cortex, from at least three brain slices cut along the A-P axis of the brain. To minimize RNA degradation, Laser Capture Microdissection was completed within 3-4 days for a single brain post cryo-sectioning.

### Sequencing library preparation for MAPseq of the OB outputs

The general procedure of RNA extraction and sequencing library preparation has been described previously^34–37^. Briefly, we extracted the total RNA of each region of interest dissected from each brain slice using RNAqueous®-Micro Total RNA Isolation Kit (Thermo Fisher Scientific) and eluted RNA in 20 µl solution. Before library preparation, we randomly selected and ran several RNA samples from each brain on Bioanalyzer (Agilent Technology) to ensure good RNA quality. We mixed 9 µl total RNA of each sample with 1 µl spike-in RNA (∼1X10^-9^ µg/µl spike-in RNAs for each target sample and 1 µl of 1X10^-7^ µg/µl spike-in RNAs for each OB sample), and reverse-transcribed the barcode mRNA into the first-strand cDNA using a gene specific barcoded reverse transcription primer and SuperScript IV (Thermo Fisher Scientific). The barcoded reverse transcription primer has sequence 5’-CTT GGC ACC CGA GAA TTC CAX XXX XXX XXX XXZ ZZZ ZZZ ZTG TAC AGC TAG CGG TGG TCG-3’, where X12 is the barcoded unique molecular identifiers (UMI) and Z8 are barcoded sample specific identifier (SSI). The synthesized 1^st^ strand cDNA of each sample was labelled with a 12 nt UMI (unique for each RNA molecule) and 8nt SSI (unique for each sample). After synthesis of the first strand cDNA, every 10 to 12 samples of target brain regions or OB (targets and OB of each brain separately before sequencing) were pooled together for clean-up by 1.8×SPRI select beads and synthesis of the second strand cDNA. The double-stranded cDNA samples were cleaned up by 1.8x SPRI beads (Beckman Coulter) and treated with Exonuclease I (New England Biolabs) to remove the single-stranded reverse transcription primers before PCR amplification.

The double-stranded cDNA samples were amplified by two rounds of nested PCR reactions using standard Accuprime Pfx protocols (Thermo Fisher Scientific) with 2 mins extension for each cycle. Primers 5’-ACG AGC TGT ACA AGT AAA CGC GT-3’ and 5’-CAA GCA GAA GAC GGC ATA CGA GAT CGT GAT GTG ACT GGA GTT CCT TGG CAC CCG AGA ATT CCA-3’ were used for the first PCR reaction and primers 5’-AAT GAT ACG GCG ACC ACC GA-3’ and 5’-CAA GCA GAA GAC GGC ATA CGA-3’ were used for the second PCR reaction. In the first PCR reaction, the pooled cDNA samples for OB target brain regions and OB were amplified for 12 cycles and 9 cycles respectively in 150 µl volume per 10-12 samples. After treatment with Exonuclease I (New England Biolabs) to remove excess primers, all the PCR1 products of target areas and OB of each brain were pooled together respectively. A quarter of the pooled PCR1 products of target areas and OB were amplified by 14-16 cycles in 8 ml and 13-15 cycles in 4 ml respectively. The PCR2 products were cleaned up and concentrated by SV wizard PCR cleanup kit (Promega), Ampure XP beads (Beckman Coulter), and finally 2% agarose gel electrophoresis. The 233 bp PCR product was cut from the agarose gel, cleaned up by Qiagen MinElute Gel Extraction Kit and quantified by Bioanalyzer (Agilent Technology) and qPCR. Afterward, we combined OB target regions and OB of two sampled brains together into one sequencing library.

### BARseq on OB slices

Mice were euthanized 24-28 hours after Sindbis virus injection by brief isofluorane anesthesia followed by decapitation, and the brains were immediately extracted. The olfactory bulb was cut off and either fixed in 4% paraformaldehyde (PFA, Electron Microscopy Sciences) in PBS, post-fixed for 24 hours, and cryoprotected before frozen, or directly frozen as previously described^36^. The bulb hemispheres were then sliced into 20 µm thick sections and processed for BARseq. BARseq library preparation and sequencing was done as previously described^36^. *In situ* sequencing was performed on either a Nikon TE-2000 or an Olympus IX-81 with a Crest X-light v2 spinning disk confocal microscope using a LDI 7-laser light source. Filters and lasers used for each imaging channel documented in **Supplementary Table 2.**

### Viral labelling of piriform cortex (PC) output neurons

Mice were anesthetized by ketamine (60 mg/kg) and xylazine (10 mg/kg) administered intraperitoneally after a brief induction with oxygenated 4% isoflurane. Mice were positioned with skull dorsal surface horizontal and mounted in a stereotaxic frame as described above and small craniotomies were performed into the skull above the piriform cortex. We injected 100 nl Sindbis viruses (1:3 diluted) unilaterally to three sites along the A-P axis of piriform cortex by positive pressure (anterior: AP +1.75 mm, ML 2.8 mm, DV 4.75 mm from the skull surface of bregma; middle: AP +0.15 mm, ML 3.9 mm, DV 5 mm from the skull surface of bregma; posterior: AP -1.5 mm, ML 4.25 mm, DV 5.5 mm from the skull surface of bregma). 24-48 hours after injection, mice were perfused with 4% PFA in PBS, and post-fixed in 4% PFA in PBS for 24 hours at 4°C, mounted into OCT in dry ice ethanol batch and stored at -80°C.

### Cryo-sectioning and dissection for MAPseq of PC outputs

The brain was sliced from the anterior to posterior end into 200 µm coronal sections with a cryostat (Leica CM 3050S). Each slice was cut by a fresh, unused part of the blade and mounted on a clean microscope glass slide, rapidly frozen on dry ice, and stored at -80° C until dissection. Brain slices were thawed, stained in 0.1 % Toluidine blue O (dissolved in 1.0 % NaCl, pH = 2.3) for 1 min, washed in PBS for 2 mins at room temperature, and kept on wet ice until dissection. Brain areas were hand-dissected from each brain slice under dissection scope by scalpel at room temperature, immediately transferred to 1.5 ml safe-lock Eppendorf tube, frozen on dry ice, and stored at -80 °C until RNA extraction. We collected the injection site, ipsilateral piriform cortex, from each brain slice individually and combined together each region of interest collected across multiple slices from a single PC target brain region. To assess cross-sample contamination, we collected as negative controls tissue from the contralateral OB and cerebellum, which are known to receive no input from the piriform cortex. Images of the brain slices were taken before and after dissection.

### Sequencing library preparation for MAPseq of PC outputs

Each dissected brain sample was digested with 8 µl of protease in 200 µl of digestion buffer from the RecoverAll^TM^ Total Nucleic Acid Isolation Kit for FFPE (Thermo Fisher Scientific). Samples containing big chunks of brain tissue were homogenized in an electronic homogenizer. Samples were then incubated at 50°C for 15 mins and then 80°C for 15 mins while shaking at 3,000 rpm. 1 ml of Trizol (Thermo Fisher Scientific) was immediately added to each digested sample and mixed well. We used half of the Trizol-sample mixture to extract the total RNA and stored the other half at -80°C.

General procedures of reverse-transcription and the rest of library preparation were performed as described above. Briefly, after synthesis of the first strand cDNA, PC target brain regions and the piriform cortex injection sites of each sampled brain were pooled together respectively for further library preparation. The double-stranded cDNA was amplified by 15 cycles for the PC target regions and 10 cycles for the piriform cortex injection sites in the first PCR reaction in 250 µl volume (per 8-10 samples) and a quarter of PCR1 products were further amplified by 6-8 cycles in 2 ml total volume for the target brain regions and 6-9 cycles in 12 ml total volume for the piriform cortex injection sites.

### Sequencing of the MAPseq samples

The pooled library constructed as described above was sequenced on an Illumina Nextseq500 high output run at paired end 36 with Illumina compatible Read 1 Sequencing Primer and Illumina compatible small RNA read 2 primer as described previously^34^.

### qPCR measurement of β-actin in OB target samples

4 µl total RNA extracted from the dissected target areas of one brain (YC61) was reverse transcribed using oligodT primers and Superscript III (Thermo Fisher Scientific) in 20 µl reaction volume. 6 µl of the reverse-transcription products was used to quantify the amount of β-actin cDNA by 3 replicate qPCR reactions in Power SYBR Green PCR Master Mix (Thermo Fisher Scientific) using primers 5’-CGG TTC CGA TGC CCT GAG GCT CTT-3’ and 5’-CGT CAC ACT TCA TGA TGG AAT TGA-3’. The relative amount of β-actin mRNA of each sample was normalized to the reference sample and calculated as 2^(CTRef – CTSample)^. Sample 7 was used as reference for normalization, which had the smallest CT value, which was consistent across 3 independent qPCR plates.

### MAPseq data processing

The raw MAPseq data consist of two fastq files, with paired end 1 covers the 30 nt barcode sequence and paired end 2 covers 12 nt UMI and the 8 nt SSI^34, 37^. The raw sequencing data was pre-processed in bash and then analyzed in Matlab (Mathworks) as previously described^34–37^. We first sorted the barcodes from each sample based on the SSI sequence and then counted the number of UMI as the molecule number of individual barcodes. Subsequently, we matched the barcodes in the injection sites to barcodes in the target brain regions based on barcode sequences and generated a barcode matrix containing the barcode sequences and their corresponding molecule counts for each sample.

We only analyzed barcodes that had molecule counts larger than 50, but smaller than 100,000 in the injection site (OB or PC) and had more molecule counts in the injection site than in any of the target brain regions sampled. To exclude low-confidence projections, we required a barcode to have at least 10 molecule counts in the PC target regions (sampled at the brain region resolution) or at least 3 molecule counts in the OB target regions (sampled at single slice level). We have also normalized the variation in library preparation and sequencing across different samples within each brain based on recovered spike-in RNA (**Supplementary Note 1**) as described previously^34, 35^.

### CAV-2 retrograde tracing to validate OB projections

Mice were anesthetized, positioned and mounted as described above in the viral labeling of PC output neurons section and small craniotomies were performed into the skull above the injection site. We first injected 6-7 weeks old C57BL/6J males (The Jackson Laboratory) unilaterally with 70 nl viral mixture of AAV pCAG-FLEX-{EGFP}_on_ (Addgene plasmid #51502, titer 1.1E+12 GC/ml,) and AAV-{CAR}_off_ (titer 1E+12 GC/ml)^104^ at 200 µm depth from the bulb surface, at three penetration sites along the A-P axis of the OB dorsal aspect. Two weeks after the initial injection, we injected 140 nl of CAV2-Cre (Montpellier vector platform, titer 2.75E+12 pp/ml) to the middle of piriform cortex (MPC, boundary between APC and PPC) or the lateral entorhinal cortex (lENT) in the same hemisphere as the initial OB injection. The stereotaxic coordinates used are the following: MPC, AP 0.10 mm, ML 3.50 mm, DV 5.50 mm and AP 0.10 mm, ML 3.85 mm, DV 5.00 mm from the skull surface of bregma; lENT, AP -3.50 mm, ML 4.40 mm, DV 4.70 mm from the skull surface of bregma. Two weeks after CAV2-Cre injection, the animals were perfused in 4% PFA in PBS and post-fixed in 4% PFA in PBS for 24 hours at 4 °C. The fixed brains were sliced into 100 µm coronal sections on a vibratome (LeicaVT1000S, Leica).

Every other brain slice containing tissue from the piriform cortex was immunostained by Goat anti-GFP antibody (Rockland Immunochemicals, Cat#600-101-215) and Donkey anti-Goat secondary antibody, Alexa Fluor 568 conjugated (Invitrogen) as described^104^. Brain slices with Cav2-Cre injection at MPC and lENT were immunostained by Rabbit anti-Cre antibody (Millipore Sigma, Cat#69050-3) and Donkey anti-Rabbit secondary antibody, Alexa Fluor 568 conjugated (Invitrogen). After immunostaining, brain slices were stained with DAPI (Sigma), mounted and imaged on an Olympus IX-81 microscope with a Crest X-light v2 spinning disk confocal microscope, an LDI 7-laser light source, and a Photometrics BSI Prime sCMOS camera. Detailed settings of lasers and filters for the imaging channels are shown in **Supplementary Table 2**.

The immunofluorescence signals of axonal branches to the piriform cortex from the labelled mitral cells were quantified using Fiji^105^. Rolling ball background subtraction was applied to remove smooth continuous background from each image. The same minimal threshold was then applied for all the images taken of the same brain and the fluorescence intensity above the chosen threshold within piriform cortex was recorded.

### Retrograde tracing to validate PC outputs

We used RetroBeads (Lumaflour) to retrogradely label PC output neurons that project to different brain regions in 8-to-10-weeks old C57BL/6J male mice (The Jackson Laboratory). Mice were anesthetized, positioned and mounted as described above and small craniotomies were performed into the skull above the injection sites. In one subset of animals (3 mice), we injected 35 nl different fluorescent microbeads (full strength green beads and 1:4 diluted read beads) into the unilateral OB and lENT respectively. In another subset of animals (2 mice), we injected 35 nl green and red beads into the unilateral AON and CoA respectively. The stereotaxic coordinates used are the following: OB, 1.20 mm anterior to the blood vessel between OB and prefrontal cortex, 0.90 mm from the middle line, 0.60 and 1.25 mm deep from the bulb surface; lENT, AP -3.50 mm, ML 4.50 mm, DV 4.75 mm from the skull surface of bregma; AON, AP 3.20 mm, ML 1.25 mm, DV 3.80 mm from the skull surface of bregma; CoA, AP -2.00 mm, ML 2.50 mm, DV 5.80 mm from the skull surface of bregma. To minimize the tracer leakage in the track, we sealed the glass pipette tip using a tiny amount of mineral oil before pipette penetration into the brain and waited for 10 mins after injection before slowly retracting the pipette.

Animals were allowed to recover for 3-4 days and perfused and fixed in 4% PFA as described above in CAV-2 retrograde tracing section. The fixed brains were sliced into 100 µm coronal sections and stained with DAPI. Every other brain slices with piriform cortex was mounted and imaged on an Olympus IX-81 microscope with a Crest X-light v2 spinning disk, an LDI 7-laser light source, and a Photometrics BSI Prime sCMOS camera. Details of lasers and filters for the imaging channels are reported in **Supplementary Table 2**.

The fluorescent signals of retrograde labelling in piriform cortex were quantified using Fiji^105^. Rolling ball background subtraction was applied to remove smooth continuous background from each image. The same minimal threshold was then applied for all the images of the same channel (red or green) of each brain. The number of pixels above the chosen threshold within piriform cortex was measured as the intensity of retrograde signals of each brain slice.

### OB projection data analysis

All analyses were based on barcode counts normalized by spike-in RNA counts. All analyses were done in MATLAB.

To examine the relationship between pMC soma positions and their projections, we calculated the “projection bias” for each A/P soma position in the bulb. To calculate the projection bias for a given A/P position, we calculated the mean projection barcode counts to an area for neurons at that A/P position. We also calculated the mean projection barcode counts after shuffling the A/P position labels of neurons. The projection bias is then defined as the ratio between the mean projection barcode counts for the experimentally observed data and that for the shuffled data, and this ratio was plotted as the relative projection strength in **Extended Data Fig. 3d,e**.

### IPR

The inverse participation ratio (IPR) is a frequently used measure of vectors’ sparseness^106^. It is related to the lifetime or population sparseness^107, 108^, however, IPR is designed to directly represents the number of non-zero components in the vector. For a vector 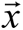 with components *x_i_* IPR is defined as 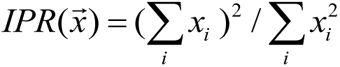. For example, for a vector which contains *N* non-zero components all equal to *c*, IPR is equal to *N*. IPR also does not depend on the scale of vector 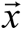 : multiplying 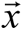 by a number, i.e. the transformation 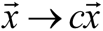, does not change its value. We therefore used IPR to quantify the average number of non-zero vector components for each barcode sequence. For the vector 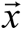 with components *x_i_* describing the number of molecules found in brain slice *i* for a particular barcode sequence, IPR gives an estimate of the number of slices substantively occupied by that barcode sequence.

### OB target region pairwise correlations

Pearson correlation between pMC projections across pairs of OB target regions was calculated in MATLAB using spike-in normalized barcode counts per brain region. Only correlations that passed statistical significance after Bonferroni correction are shown in **Fig. 1k**.

### Classifiers

To cluster the MAPseq barcoded neurons into pMCs, pTCs and pDC, we used a three-layer fully connected feedforward neural network (NN). In the BARseq data set, barcoded neurons of different cell types were clearly identified based on their positions within the olfactory bulb (distance from surface and from the mitral cell layer respectively). We then trained a NN to predict the cell types using the projections of the BARseq identified neurons to the six OB target regions sampled (265/415 neurons sampled via BARseq). The NN was further used to estimate the probability of assigning a barcoded neuron to a certain type using the MAPseq projection dataset (6 brains). We used the trained NN to obtain the classification probabilities of 4,894 barcoded neurons from the MAPseq dataset. A 0.85 or higher probability of belonging to the pMC cluster was considered a hit, and the corresponding barcoded neuron labeled as pMC. Details of the classifier are described in **Supplementary Note 2**.

### Conditional probability analysis

We used conditional probability as a measure of normalized co-innervation strength between different locations in the piriform cortex (PC) and target regions (AON, OT, CoA, lENT). The conditional probability is interpreted as the probability of observing a barcode molecule within the target regions, given that a molecule of the same barcode was also observed in a specific slice in PC. Our calculation is based on the matrix *C*(*b*, *s*) of spike-in normalized molecule counts for the barcode *b* in piriform cortex slice *s* from the MAPseq data. This matrix was used to calculate the probability of finding a barcode molecule in a given PC slice. Across the PC slices, this probability is 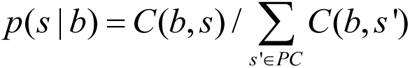 and across the target regions, enumerated by the index *r* = 1..4, it is 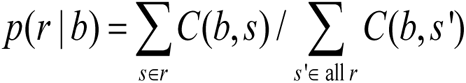. These two conditional probabilities formed the basis for our calculation.

Since most OB pMC project to PC, we can assume that these barcodes are present in PC. We then reasoned that the probabilities of these barcoded neurons to be present in PC should therefore be equal, i.e. *p*(*b*) = 1/ *N_B_*, where *N_B_* is the total number of barcodes detected in PC. We have verified that our conclusions are not affected by this assumption. Indeed, adopting a different model for *p*(*b*), such as 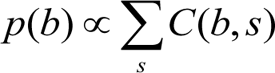 does not noticeably affect our results. We further used the law of total probability to calculate the probability of a barcode molecule being sampled from a given slice in PC: *p*(*s*) = ∑*_b_ p*(*s* | *b*) *p*(*b*).

To compute the conditional probability of finding a barcode molecule *b* in a given PC slice *s*, we used Bayes’ theorem: *p*(*b* | *s*) = *p*(*s* | *b*) *p*(*b*) / *p*(*s*). Then, using the law of total probability, we calculated the probability that a barcode molecule comes from a given target region, granted that another molecule of the same barcode is found in PC slice *s* : *p*(*r* | *s*) = ∑*_b_ p*(*r* | *b*) *p*(*b* | *s*). Simplifying the calculation under the uniform sampling of barcoded neurons *p*(*b*) = const, we calculated the conditional probability of projecting to a given target region given that molecules of the same barcode are found in the PC slice:

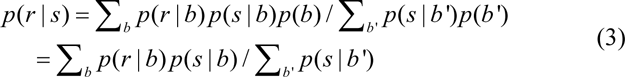

### Regression analysis and shuffled controls

To determine the robustness of our analysis to sampling effects, we used bootstrap to re-sample the set of barcodes with replacements and recalculate the conditional probability matrices. Piecewise linear regressions on APC and PPC vs. single linear regression along the whole of PC anterior-posterior axis better fitted the data as indicated by using 10 fold cross-validation (**Supplementary Table 3**). Similar results were obtained shuffling the PC slices by randomly permuting the AP positioning order in the conditional probability matrix (data not shown).

### Down-sampling controls

To investigate the effects of lower throughput on reaching statistical significance on our observations, we down-sampled the pMC neurons of our MAPseq dataset to various sizes. On these sets, we calculated the conditional probability matrices and performed the same analyses. We note that most of our results become statistically insignificant on these smaller datasets, suggesting that large sample size is necessary to discover the patterns of projection observed in our data (**Extended Data Fig. 4d**).

### Tiling

To quantify tiling, we sorted the barcoded neurons according to the A-P location of the maximum projection count in different slices (**Fig. 2a**, **f**, **g**, etc.). To separate the barcoded neurons into narrowly and broadly projecting populations, we computed their IPR and average A-P PC locations, and build the 2D distribution according to these quantities (**Fig. 2d**). Narrowly and broadly projecting groups could be well-separated using a watershed algorithm in the 2D plane. The boundary generated by the watershed algorithm in shown in **Fig. 2d** by the white line. We performed exponential fits (red curves in **Fig. 2**) using standard non-linear fitting functions available in MATLAB (Mathworks). The functional model used was *x*(*r*) = *w* exp(−*α r*), where *x*, *r*, *w*, and *α* are the A-P location of a barcoded neuron projection maximum, A-P rank of the barcoded cell (neuron number), the width of the tiled brain region corrected for edge effects (number of slices - 1), and the only fitting parameter, representing the exponent’s decay.

### Piriform cortex projection data analysis

For all analysis of intra-PC projections, we excluded the three slices adjacent to each soma location (one slice on each side and the central slice, representing ∼600 µm along the A-P axis) because of possible contribution of barcode molecules from the dendrites of the barcoded neurons. To cluster PC slices, we first re-sampled with replacement 500 neurons for each A-P position to equalize the number of neurons from different A-P positions, then used the correlation of barcode molecule counts between PC slices to build an adjacency matrix and performed Louvain community detection^109^. To compare PC outputs to the Allen projection atlas, we calculated the Pearson correlation between the mean barcode molecule counts for each area and GFP intensities from the Allen projection atlas (**Extended Data Fig. 7**). Neurons in APC were compared to experiment ID 127907465 and neurons in PPC were compared to experiment ID 146857301.

We fitted the piriform projections using the Lorentzian function *f* (*x*) = *A* / (1+ *x*^2^ / *w*^2^). Here *A* and *w* are the fitting parameters representing the amplitude and the width of the function respectively. These parameters were found using standard non-linear fit function of MATLAB (Mathworks, Inc.). The values of the twice the distribution width are marked in **Fig. 3d**. Thus, for intra PC connectivity, we obtain *w* ≈ 0.5mm, leading to the curve’s width at half height of 1mm as indicated in **Fig. 3d**.

To identify groups of PC neurons based on output projections, we performed two layers of Louvain community detection, then manually combined clusters resulted from the second layer if they were intermingled in the t-sne plots (**Fig. 3e**). To determine the relative anterior/posterior bias of intra-piriform projections, for each neuron in slice X along the A-P axis of the piriform cortex, we calculated the difference in barcode molecule counts in slices X - 8 to X - 3 (anterior to the soma location X) and barcode molecule counts in slices X + 3 to X + 8 (posterior to the soma location), normalized by the total barcode molecule counts in all these slices. Neurons that did not have eight slices on either side (i.e. neurons in the first eight and last eight slices in the PC) were excluded from this analysis; p values were calculated using a sign rank test (**Extended Data Fig. 8f**).

To predict soma positions using the piriform output projections, we trained a linear regression model using the piriform output projections as inputs to predict soma A/P positions. We also trained a random forest model (default settings using TreeBagger in MATLAB with 200 trees) in regression mode to predict soma A/P positions using the group labels. For both models, the performance was evaluated using 10-fold cross validation (**Extended Data Fig. 8d**,**e**).

### Matching of PC input and output circuit motifs

To check for input-output circuit motifs, we calculated the conditional probability matrix from the PC injection data, that is the probability of a barcode molecule being sampled from a target region, given that the barcoded neuronal somata is located in a given PC slice. The PC output projection data on target regions was pooled over each target region (*r*) sampled. To compute the conditional probability of projecting from PC to another region, *p_PCx_* (*r* | *s*), the equations described above [Equation (3)] were slightly amended. Indeed, *p_PCx_* (*r* | *s*) reflects the rate of projections given that the soma is present at slice *s*, not an axonal collateral/branch, as in *p*(*r* | *s*) [Equation (3)].

Instead of using the distribution of molecules of a barcode 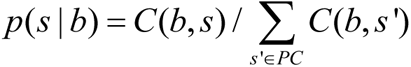, we used a delta function as the probability distribution of the somata labeled by that barcode: *p*_soma_ (*s* | *b*) = *δ* (*s*, arg max(*B*(*s*’, *b*))), where *B*(*s*, *b*) are the spike-in normalized molecule count of barcode *b* in a PC viral injection experiment. We then repeated the calculations that led to Equation (3) to compute *p_PCx_* (*r* | *s*) (**Fig. 4a**). The probability of barcode *b* was calculated as *p*(*r* | *b*) = *C*(*r*, *b*) (∑_*r*∈*t*arg*et*_ *C*(*r*, *b*)),the number of barcodes with where *C*(*r*,*b*) is the counts of the barcode measured in target region *r*. We evaluated 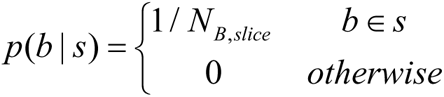 directly, where *N_B,slice_* is the number of barcodes with somas located at the specified PC slice. The conditional probability from these quantities was calculated as *p*(*r* | *s*) = ∑*_b_ p*(*r* | *b*) *p*(*b* | *s*). This conditional probability matrix is interpreted as the mean normalized projection strength from PC to a given target region. We further performed PCA on the N × 4 projection matrix containing the projection strengths to the four extra-piriform areas (AON, OT, CoA, and lENT), where N is the number of PC output neurons. The mean loadings of the first two principal components for PC neurons at each A-P location were then plotted to examine the slope change in projection gradients at the APC/PPC boundary (**Fig. 4b**).

We further calculated the Spearman correlation between the conditional probability matrix of PC inputs and mean projection strength matric of PC outputs along the A-P axis of piriform cortex with respect to the four extra-piriform target regions sampled (**Fig. 4c**). To visualize the evolution of the projection strengths along the AP axis of the PC, we used a Bézier curve to interpolate a curve representing the two conditional probabilities. We used the formula 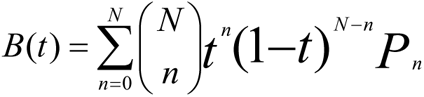; where *P_n_* is the coordinate constructed by the PC slice *n* +1 and the entries in the two conditional probability matrices.

## Data availability

The raw sequencing data for all *in vitro* high-throughput sequencing reported in this study (including six OB MAPseq brains, YC61, YC65, YC86, YC92, YC109, and YC111, two BARseq brains, YC107 and YC113, and five PC MAPseq brains, XC113, XC119, XC120, XC127, and YC123 are available on SRA with accession number PRJNA707572. Images for MAPseq brain dissection and a list of Allen Brain Reference Atlas coronal levels corresponding to the dissected brain slices are available on Mendeley Data (https://data.mendeley.com/datasets/ggbft4btrb/draft?a=8032c065-dc84-4906-b5ce-445902116f7a). All data matrices representing OB and PC output neuron identities, soma locations and projection patterns included in the analyses presented here are available on Mendeley Data (https://data.mendeley.com/datasets/6t9mb3yydy/draft?a=8a33a230-3176-40ca-9ad9-a8727afa31b6).

## Code availability

The code used for analysis is available on Mendeley Data (https://data.mendeley.com/datasets/6t9mb3yydy/draft?a=8a33a230-3176-40ca-9ad9-a8727afa31b6) and on Github (https://github.com/KoulakovLab/OlfactoryProjectomeAnalysis).

## Acknowledgments

The authors would like to acknowledge A. Banerjee, K.M. Franks, D. Fürth, P. Gupta, S. Navlakha, G.B. Keller, S. Lu, F. Marbach, X. Zheng and members of the Albeanu, Koulakov and Zador groups for support and critical discussions. This work was supported by the following funding sources: BRAIN 1R01NS111673 to D.F.A. and A.A.K. and TR01 5R01DC017876 to D.F.A and A.A.K.

## References

1. Kaas, J. H. Topographic maps are fundamental to sensory processing. Brain Res. Bull. 44, 107–112 (1997).

2. Knudsen, E. I., du Lac, S. & Esterly, S. D. Computational maps in the brain. Annu. Rev. Neurosci. 10, 41–65 (1987).

3. Chklovskii, D. B. & Koulakov, A. A. Maps in the brain: what can we learn from them? Annu. Rev. Neurosci. 27, 369–392 (2004).

4. Brewer, A. A. & Barton, B. Maps of the Auditory Cortex. Annu. Rev. Neurosci. 39, 385–407 (2016).

5. Harding-Forrester, S. & Feldman, D. E. Somatosensory maps. Handb. Clin. Neurol. 151, 73–102 (2018).

6. Nauhaus, I. & Nielsen, K. J. Building maps from maps in primary visual cortex. Curr. Opin. Neurobiol. 24, 1–6 (2014).

7. Price, J. L. An autoradiographic study of complementary laminar patterns of termination of afferent fibers to the olfactory cortex. J. Comp. Neurol. 150, 87–108 (1973).

8. Haberly, L. B. & Price, J. L. The axonal projection patterns of the mitral and tufted cells of the olfactory bulb in the rat. Brain Res. 129, 152–157 (1977).

9. Skeen, L. C. & Hall, W. C. Efferent projections of the main and the accessory olfactory bulb in the tree shrew (Tupaia glis). J. Comp. Neurol. 172, 1–35 (1977).

10. Luskin, M. B. & Price, J. L. The distribution of axon collaterals from the olfactory bulb and the nucleus of the horizontal limb of the diagonal band to the olfactory cortex, demonstrated by double retrograde labeling techniques. J. Comp. Neurol. 209, 249–263 (1982).

11. Haberly, L. B. Parallel-distributed processing in olfactory cortex: new insights from morphological and physiological analysis of neuronal circuitry. Chem. Senses 26, 551–576 (2001).

12. Gottfried, J. A. Central mechanisms of odour object perception. Nat. Rev. Neurosci. 11, 628–641 (2010).

13. Nagayama, S. et al. Differential axonal projection of mitral and tufted cells in the mouse main olfactory system. Front. Neural Circuits 4, (2010).

14. Igarashi, K. M. et al. Parallel mitral and tufted cell pathways route distinct odor information to different targets in the olfactory cortex. J. Neurosci. Off. J. Soc. Neurosci. 32, 7970–7985 (2012).

15. Sosulski, D. L., Bloom, M. L., Cutforth, T., Axel, R. & Datta, S. R. Distinct representations of olfactory information in different cortical centres. Nature 472, 213–216 (2011).

16. Ghosh, S. et al. Sensory maps in the olfactory cortex defined by long-range viral tracing of single neurons. Nature 472, 217–220 (2011).

17. Miyamichi, K. et al. Cortical representations of olfactory input by trans-synaptic tracing. Nature 472, 191–196 (2011).

18. Bekkers, J. M. & Suzuki, N. Neurons and circuits for odor processing in the piriform cortex. Trends Neurosci. 36, 429–438 (2013).

19. Choi, G. B. et al. Driving opposing behaviors with ensembles of piriform neurons. Cell 146, 1004– 1015 (2011).

20. Wilson, D. A. & Sullivan, R. M. Cortical processing of odor objects. Neuron 72, 506–519 (2011).

21. Davison, I. G. & Ehlers, M. D. Neural circuit mechanisms for pattern detection and feature combination in olfactory cortex. Neuron 70, 82–94 (2011).

22. Giessel, A. J. & Datta, S. R. Olfactory maps, circuits and computations. Curr. Opin. Neurobiol. 24, 120–132 (2014).

23. Stettler, D. D. & Axel, R. Representations of odor in the piriform cortex. Neuron 63, 854–864 (2009).

24. Johnson, D. M. G., Illig, K. R., Behan, M. & Haberly, L. B. New Features of Connectivity in Piriform Cortex Visualized by Intracellular Injection of Pyramidal Cells Suggest that “Primary” Olfactory Cortex Functions Like “Association” Cortex in Other Sensory Systems. J. Neurosci. 20, 6974–6982 (2000).

25. Franks, K. M. et al. Recurrent circuitry dynamically shapes the activation of piriform cortex. Neuron 72, 49–56 (2011).

26. Babadi, B. & Sompolinsky, H. Sparseness and expansion in sensory representations. Neuron 83, 1213–1226 (2014).

27. Barak, O., Rigotti, M. & Fusi, S. The sparseness of mixed selectivity neurons controls the generalization-discrimination trade-off. J. Neurosci. Off. J. Soc. Neurosci. 33, 3844–3856 (2013).

28. Rigotti, M. et al. The importance of mixed selectivity in complex cognitive tasks. Nature 497, 585– 590 (2013).

29. Hansel, D. & van Vreeswijk, C. The mechanism of orientation selectivity in primary visual cortex without a functional map. J. Neurosci. Off. J. Soc. Neurosci. 32, 4049–4064 (2012).

30. Schaffer, E. S. et al. Odor Perception on the Two Sides of the Brain: Consistency Despite Randomness. Neuron 98, 736–742.e3 (2018).

31. Litwin-Kumar, A., Harris, K. D., Axel, R., Sompolinsky, H. & Abbott, L. F. Optimal Degrees of Synaptic Connectivity. Neuron 93, 1153–1164.e7 (2017).

32. Krishnamurthy, K., Hermundstad, A. M., Mora, T., Walczak, A. M. & Balasubramanian, V. Disorder and the neural representation of complex odors: smelling in the real world. bioRxiv 160382 (2017) doi:10.1101/160382.

33. Stevens, C. F. What the fly’s nose tells the fly’s brain. Proc. Natl. Acad. Sci. 112, 9460–9465 (2015).

34. Kebschull, J. M. et al. High-Throughput Mapping of Single-Neuron Projections by Sequencing of Barcoded RNA. Neuron 91, 975–987 (2016).

35. Han, Y. et al. The logic of single-cell projections from visual cortex. Nature 556, 51–56 (2018).

36. Chen, X. et al. High-Throughput Mapping of Long-Range Neuronal Projection Using In Situ Sequencing. Cell 179, 772–786.e19 (2019).

37. Huang, L. et al. BRICseq Bridges Brain-wide Interregional Connectivity to Neural Activity and Gene Expression in Single Animals. Cell 182, 177–188.e27 (2020).

38. Sun, Y.-C. et al. Integrating barcoded neuroanatomy with spatial transcriptional profiling reveals cadherin correlates of projections shared across the cortex. bioRxiv 2020.08.25.266460 (2020) doi:10.1101/2020.08.25.266460.

39. Mombaerts, P. Axonal wiring in the mouse olfactory system. Annu. Rev. Cell Dev. Biol. 22, 713– 737 (2006).

40. Wilson, R. I. & Mainen, Z. F. Early events in olfactory processing. Annu. Rev. Neurosci. 29, 163– 201 (2006).

41. Soucy, E. R., Albeanu, D. F., Fantana, A. L., Murthy, V. N. & Meister, M. Precision and diversity in an odor map on the olfactory bulb. Nat Neurosci 12, 210–220 (2009).

42. Boyd, A. M., Sturgill, J. F., Poo, C. & Isaacson, J. S. Cortical feedback control of olfactory bulb circuits. Neuron 76, 1161–1174 (2012).

43. Otazu, G. H., Chae, H., Davis, M. B. & Albeanu, D. F. Cortical Feedback Decorrelates Olfactory Bulb Output in Awake Mice. Neuron 86, 1461–1477 (2015).

44. Boyd, A. M., Kato, H. K., Komiyama, T. & Isaacson, J. S. Broadcasting of cortical activity to the olfactory bulb. Cell Rep. 10, 1032–1039 (2015).

45. Markopoulos, F., Rokni, D., Gire, D. H. & Murthy, V. N. Functional properties of cortical feedback projections to the olfactory bulb. Neuron 76, 1175–1188 (2012).

46. Oswald, A.-M. & Urban, N. N. There and Back Again: The Corticobulbar Loop. Neuron 76, 1045– 1047 (2012).

47. Scott, J. W., McBride, R. L. & Schneider, S. P. The organization of projections from the olfactory bulb to the piriform cortex and olfactory tubercle in the rat. J. Comp. Neurol. 194, 519–534 (1980).

48. Murthy, V. N. Olfactory Maps in the Brain. Annu. Rev. Neurosci. 34, 233–258 (2011).

49. Hagiwara, A., Pal, S. K., Sato, T. F., Wienisch, M. & Murthy, V. N. Optophysiological analysis of associational circuits in the olfactory cortex. Front. Neural Circuits 6, (2012).

50. Bolding, K. A. & Franks, K. M. Recurrent cortical circuits implement concentration-invariant odor coding. Science 361, (2018).

51. Pashkovski, S. L. et al. Structure and flexibility in cortical representations of odour space. Nature 583, 253–258 (2020).

52. Caron, S. J. C., Ruta, V., Abbott, L. F. & Axel, R. Random convergence of olfactory inputs in the Drosophila mushroom body. Nature 497, 113–117 (2013).

53. Grimaud, J., Dorrell, W., Pehlevan, C. & Murthy, V. Bilateral Alignment of Receptive Fields in the Olfactory Cortex Points to Non-Random Connectivity. bioRxiv 2020.02.24.960922 (2020) doi:10.1101/2020.02.24.960922.

54. Zheng, Z. et al. Structured sampling of olfactory input by the fly mushroom body. bioRxiv 2020.04.17.047167 (2020) doi:10.1101/2020.04.17.047167.

55. Gruntman, E. & Turner, G. C. Integration of the olfactory code across dendritic claws of single mushroom body neurons. Nat. Neurosci. 16, 1821–1829 (2013).

56. Zavitz, D., Amematsro, E. A., Borisyuk, A. & Caron, S. J. C. Connectivity patterns shape sensory representation in a cerebellum-like network. bioRxiv 2021.02.10.430647 (2021) doi:10.1101/2021.02.10.430647.

57. Li, F. et al. The connectome of the adult Drosophila mushroom body provides insights into function. eLife 9, e62576 (2020).

58. Eichler, K. et al. The complete connectome of a learning and memory centre in an insect brain. Nature 548, 175–182 (2017).

59. Dasgupta, S., Stevens, C. F. & Navlakha, S. A neural algorithm for a fundamental computing problem. Science 358, 793–796 (2017).

60. Srinivasan, S. & Stevens, C. F. Scaling Principles of Distributed Circuits. Curr. Biol. 29, 2533–2540.e7 (2019).

61. Peng, H. et al. Brain-wide single neuron reconstruction reveals morphological diversity in molecularly defined striatal, thalamic, cortical and claustral neuron types. bioRxiv 675280 (2020) doi:10.1101/675280.

62. Winnubst, J. et al. Reconstruction of 1,000 Projection Neurons Reveals New Cell Types and Organization of Long-Range Connectivity in the Mouse Brain. Cell 179, 268–281.e13 (2019).

63. Oettl, L.-L. et al. Oxytocin Enhances Social Recognition by Modulating Cortical Control of Early Olfactory Processing. Neuron 90, 609–621 (2016).

64. Kikuta, S. et al. Neurons in the anterior olfactory nucleus pars externa detect right or left localization of odor sources. Proc. Natl. Acad. Sci. 107, 12363–12368 (2010).

65. Esquivelzeta Rabell, J., Mutlu, K., Noutel, J., Martin del Olmo, P. & Haesler, S. Spontaneous Rapid Odor Source Localization Behavior Requires Interhemispheric Communication. Curr. Biol. 27, 1542–1548.e4 (2017).

66. Wang, C. Y., Liu, Z., Ng, Y. H. & Südhof, T. C. A Synaptic Circuit Required for Acquisition but Not Recall of Social Transmission of Food Preference. Neuron 107, 144–157.e4 (2020).

67. Wang, C. et al. Egocentric coding of external items in the lateral entorhinal cortex. Science 362, 945–949 (2018).

68. Root, C. M., Denny, C. A., Hen, R. & Axel, R. The participation of cortical amygdala in innate, odour-driven behaviour. Nature 515, 269–273 (2014).

69. Tsao, A. et al. Integrating time from experience in the lateral entorhinal cortex. Nature 561, 57–62 (2018).

70. Fukunaga, I., Berning, M., Kollo, M., Schmaltz, A. & Schaefer, A. T. Two distinct channels of olfactory bulb output. Neuron 75, 320–329 (2012).

71. Jordan, R., Fukunaga, I., Kollo, M. & Schaefer, A. T. Active Sampling State Dynamically Enhances Olfactory Bulb Odor Representation. Neuron 98, 1214–1228.5 (2018).

72. Kapoor, V., Provost, A. C., Agarwal, P. & Murthy, V. N. Activation of raphe nuclei triggers rapid and distinct effects on parallel olfactory bulb output channels. Nat. Neurosci. 19, 271–282 (2016).

73. Yamada, Y. et al. Context- and Output Layer-Dependent Long-Term Ensemble Plasticity in a Sensory Circuit. Neuron 93, 1198–1212.e5 (2017).

74. Burton, S. D. & Urban, N. N. Greater excitability and firing irregularity of tufted cells underlies distinct afferent-evoked activity of olfactory bulb mitral and tufted cells. J. Physiol. 592, 2097–2118 (2014).

75. Geramita, M. & Urban, N. N. Differences in Glomerular-Layer-Mediated Feedforward Inhibition onto Mitral and Tufted Cells Lead to Distinct Modes of Intensity Coding. J. Neurosci. Off. J. Soc. Neurosci. 37, 1428–1438 (2017).

76. Geramita, M. A., Burton, S. D. & Urban, N. N. Distinct lateral inhibitory circuits drive parallel processing of sensory information in the mammalian olfactory bulb. eLife 5, (2016).

77. Gire, D. H. et al. Mitral cells in the olfactory bulb are mainly excited through a multistep signaling path. J. Neurosci. Off. J. Soc. Neurosci. 32, 2964–2975 (2012).

78. Eyre, M. D., Antal, M. & Nusser, Z. Distinct deep short-axon cell subtypes of the main olfactory bulb provide novel intrabulbar and extrabulbar GABAergic connections. J. Neurosci. Off. J. Soc. Neurosci. 28, 8217–8229 (2008).

79. Kosaka, T. & Kosaka, K. Heterogeneity of nitric oxide synthase-containing neurons in the mouse main olfactory bulb. Neurosci. Res. 57, 165–178 (2007).

80. Chklovskii, D. B., Schikorski, T. & Stevens, C. F. Wiring Optimization in Cortical Circuits. Neuron 34, 341–347 (2002).

81. Stevens, C. F. 1.17 - Principles of Brain Scaling. in Evolution of Nervous Systems (ed. Kaas, J. H.) 273–282 (Academic Press, 2007). doi:10.1016/B0-12-370878-8/00112-9.

82. Wang, X.-J. & Kennedy, H. Brain structure and dynamics across scales: In search of rules. Curr. Opin. Neurobiol. 37, 92–98 (2016).

83. Rodieck, R. W. The First Steps in Seeing. (Sinauer Associates is an imprint of Oxford University Press, 1998).

84. Luna, V. M. & Pettit, D. L. Asymmetric rostro-caudal inhibition in the primary olfactory cortex. Nat. Neurosci. 13, 533–535 (2010).

85. Large, A. M. et al. Differential inhibition of pyramidal cells and inhibitory interneurons along the rostrocaudal axis of anterior piriform cortex. Proc. Natl. Acad. Sci. 115, E8067–E8076 (2018).

86. Chen, C.-F. F. et al. Nonsensory target-dependent organization of piriform cortex. Proc. Natl. Acad. Sci. U. S. A. 111, 16931–16936 (2014).

87. Datiche, F., Litaudon, P. & Cattarelli, M. Intrinsic association fiber system of the piriform cortex: A quantitative study based on a cholera toxin B subunit tracing in the rat. J. Comp. Neurol. 376, 265– 277 (1996).

88. Markov, N. T. et al. Weight consistency specifies regularities of macaque cortical networks. Cereb. Cortex N. Y. N 1991 21, 1254–1272 (2011).

89. Song, S., Sjöström, P. J., Reigl, M., Nelson, S. & Chklovskii, D. B. Highly nonrandom features of synaptic connectivity in local cortical circuits. PLoS Biol. 3, e68 (2005).

90. Zeppilli, S. et al. Molecular characterization of projection neuron subtypes in the mouse olfactory bulb. bioRxiv 2020.11.30.405571 (2020) doi:10.1101/2020.11.30.405571.

91. Mazo, C., Grimaud, J., Shima, Y., Murthy, V. N. & Lau, C. G. Distinct projection patterns of different classes of layer 2 principal neurons in the olfactory cortex. Sci. Rep. 7, 8282 (2017).

92. Diodato, A. et al. Molecular signatures of neural connectivity in the olfactory cortex. Nat. Commun. 7, 12238 (2016).

93. Klyachko, V. A. & Stevens, C. F. Connectivity optimization and the positioning of cortical areas. Proc. Natl. Acad. Sci. 100, 7937–7941 (2003).

94. Barnes, D. C., Hofacer, R. D., Zaman, A. R., Rennaker, R. L. & Wilson, D. A. Olfactory perceptual stability and discrimination. Nat. Neurosci. 11, 1378–1380 (2008).

95. He, K., Zhang, X., Ren, S. & Sun, J. Deep Residual Learning for Image Recognition. in 2016 IEEE Conference on Computer Vision and Pattern Recognition (CVPR) 770–778 (IEEE, 2016). doi:10.1109/CVPR.2016.90.

96. Wang, Q. & Burkhalter, A. Stream-related preferences of inputs to the superior colliculus from areas of dorsal and ventral streams of mouse visual cortex. J. Neurosci. Off. J. Soc. Neurosci. 33, 1696– 1705 (2013).

97. Cang, J. & Feldheim, D. A. Developmental Mechanisms of Topographic Map Formation and Alignment. Annu. Rev. Neurosci. 36, 51–77 (2013).

98. Tikidji-Hamburyan, R. A., El-Ghazawi, T. A. & Triplett, J. W. Novel Models of Visual Topographic Map Alignment in the Superior Colliculus. PLOS Comput. Biol. 12, e1005315 (2016).

99. LeDoux, J. E., Farb, C. R. & Romanski, L. M. Overlapping projections to the amygdala and striatum from auditory processing areas of the thalamus and cortex. Neurosci. Lett. 134, 139–144 (1991).

100. Ledoux, J. The Emotional Brain: The Mysterious Underpinnings of Emotional Life. (Simon and Schuster, 1998).

101. van Strien, N. M., Cappaert, N. L. M. & Witter, M. P. The anatomy of memory: an interactive overview of the parahippocampal-hippocampal network. Nat. Rev. Neurosci. 10, 272–282 (2009).

102. Basu, J. & Siegelbaum, S. A. The Corticohippocampal Circuit, Synaptic Plasticity, and Memory. Cold Spring Harb. Perspect. Biol. 7, (2015).

103. Keller, G. B. & Mrsic-Flogel, T. D. Predictive Processing: A Canonical Cortical Computation. Neuron 100, 424–435 (2018).

104. Li, S.-J., Vaughan, A., Sturgill, J. F. & Kepecs, A. A Viral Receptor Complementation Strategy to Overcome CAV-2 Tropism for Efficient Retrograde Targeting of Neurons. Neuron 98, 905–917.e5 (2018).

105. Schindelin, J. et al. Fiji: an open-source platform for biological-image analysis. Nat. Methods 9, 676–682 (2012).

106. Wegner, F. Inverse participation ratio in 2+ε dimensions. Z. Für Phys. B Condens. Matter 36, 209–214 (1980).

107. Willmore, B. & Tolhurst, D. J. Characterizing the sparseness of neural codes. Netw. Bristol Engl. 12, 255–270 (2001).

108. Treves, A. & Rolls, E. T. What determines the capacity of autoassociative memories in the brain? Netw. Comput. Neural Syst. 2, 371–397 (1991).

109. Blondel, V. D., Guillaume, J.-L., Lambiotte, R. & Lefebvre, E. Fast unfolding of communities in large networks. J. Stat. Mech. Theory Exp. 2008, P10008 (2008).

