## Supplementary Note for "Wiring logic of the early rodent olfactory system revealed by high-throughput sequencing of single neuron projections"

### Supplementary Information

#### Supplementary Notes

##### Supplementary Note 1: Mapping the brain-wide projections of individual olfactory bulb output neurons via MAPseq

Multiplexed Analysis of Projections by sequencing (MAPseq) uses Sindbis virus libraries[34–38] to introduce barcoded RNA into cells in 'source' brain areas of interest (e.g. olfactory bulb, OB, **Fig. 1a**). The extent of the MAPseq projection data depends on the efficiency of neuronal infection by the virus. To label mitral and tufted cells in the olfactory bulb, we injected barcoded Sindbis virus at multiple sites along the anterior-posterior (A-P) axis on both the dorsal and ventral aspects of the bulb in one hemisphere per experiment (**Extended Data Fig. 1a**). On average, this approached labeled ~800 mitral, tufted and deep cells per bulb hemisphere, amounting to 4,894 olfactory bulb projection neurons recovered from six brains.

The spatial resolution of the MAPseq projection data depends on the parameters used for tissue dissection. Since the entire piriform cortex (PC) covers a long (~5 mm) and narrow (~1-2 mm) portion on the ventral-lateral aspect of the mouse brain, we focused on investigating the projection patterns of bulb output neurons along the A-P axis of piriform cortex. To achieve mapping at fine resolution, we cut 200 µm coronal sections along the A-P axis of the brain (**Fig. 1a**). We sampled all 6 major olfactory bulb target brain regions: anterior olfactory nucleus (AON), piriform cortex (anterior, APC and posterior, PPC), olfactory tubercle (OT), cortical amygdala (CoA), and lateral entorhinal cortex (lENT). From each coronal section, we dissected different regions of interest using laser capture micro-dissection (LCM, **Fig. 1c**); to minimize contamination from fibers of passage, we avoided the lateral olfactory tract (LOT). We also sampled the hippocampus in two brains, but found very few barcodes in this area, indicative of weak olfactory bulb projections. We thus did not harvest it in the other four brains and focused our analyses on the six target regions mentioned above. The sampled olfactory bulb target regions could be readily separated from each other by LCM based on location and Nissl staining. To minimize cross-contamination between these olfactory bulb target regions, we avoided harvesting tissue at the inter-region boundaries. For example, for each sampled bulb target region, we discarded 1-2 tissue slices collected from the most anterior and posterior ends, and also skipped a thin slab of tissue (<0.1 mm wide, **Fig. 1c**) between the anterior piriform cortex and olfactory tubercle.

Images acquired after LCM for each of the brain slices sampled are available on Mendeley Data: https://data.mendeley.com/datasets/ggbft4btrb/draft?a=8032c065-dc84-4906-b5ce-445902116f7a.

To align the A-P positions of olfactory bulb projections across different brains, we registered each coronal slice to the Allen Brain atlas. We first registered several slices using landmarks such as the closure of corpus callosum and anterior commissure, as well as the anterior end of the hippocampus. Subsequently, we matched the rest of the slices based on the ~200 µm nominal distance between two adjacent slices. After registration, we only kept samples that were common across different brains and discarded extra samples, which usually were collected at the beginning or the end of a given brain region and received no, or minimal olfactory bulb input. Our registration aligns well with the non-rigid registration generated by the whole brain software (http://www.wholebrainsoftware.org/)[110]. The average difference between these two registration methods for individual slices was 1.4 ± 1.2 (mean ± std) coronal levels, equivalent to 140 ± 120 µm. The differences in registration across the two methods used may arise from imperfect Nissl staining and cryo-sectioning, especially near the anterior and posterior ends of a brain.

The false positive rate of MAPseq in the data collected was low: only 1.74% barcodes could be detected (have non-zero barcode counts) in negative control samples known to receive no input from the olfactory bulb. Importantly, for these barcodes which had non-zero counts, the mean molecule count was 2 compared to 203 in an olfactory bulb target brain region.

We normalized the variation in library preparation and sequencing across different samples for each brain based on the recovered spike-in RNA added to each sample. To examine whether variations in area size or tissue thickness could contribute to differences in barcode counts, we compared this normalization method to normalization by: 1) dissected area size and 2) tissue volume (measured by qPCR of β-actin) for one brain (YC61). Normalization based on spike-in RNA alone, normalization based on both spike-in RNA and area size, or on tissue volume produced similar projection patterns (**Extended Data Fig.1b**) and did not change the correlations in olfactory bulb projections to piriform cortex and extra-piriform areas reported (**Figs. 1**, **2**, **Extended Data Figs. 1b-d**). We therefore normalized only by spike-in RNA for the other brains sampled as also done in previous MAPseq and BARseq studies[34–38].

**Supplementary Note 2: Classification of the olfactory bulb projection neurons**

We first trained a neural network-based classifier to distinguish mitral, tufted and deep cells based on the brain region-level projection patterns of the BARseq sampled neurons. To train this classifier, we defined a set of template neurons based on the locations of their cell bodies: putative tufted cells (pTC) were defined as those olfactory bulb projection neurons whose somata were at least 150 μm away from the mitral cell layer (MCL), and less than 100 μm from the inner surface of glomeruli; putative mitral cells (pMC) as those bulb projection neurons whose somata lied within 50 μm of the mitral cell layer; finally, putative deep cells (pDC) were defined as those with somata located 100 μm below MCL, or deeper (**Fig. 1f**). Multiple types of tufted cells exist across different layers in the olfactory bulb, including some very close, or embedded within MCL[111]. As such, pTC used as templates comprised largely of tufted cells located more superficially, whereas pMC neurons may represent a mixture of mostly mitral cells, as well as a minority of deep tufted cells displaced in the MCL.

The classifier was based on a two-layer feed-forward neural network that computed the Bayesian probability of a neuron to belong to different cell types (pMCs/pTCs/pDCs) given their density of projection to the six olfactory bulb target regions sampled. The classifier was optimized so as to minimize classification error (softmax) in the BARseq dataset (265 mitral cells/tufted cells/deep cells defined based on soma position in the olfactory bulb), and did not distinguish barcodes based on their fine distributions of projection within each target brain region (**Fig. 1f; Extended Data Fig. 2a**). Our classifier identified pMC with high accuracy (TPR = 94%, FPR = 10%), but performed worse for identifying pDC and pTC (**Fig. 1f**). The higher performance at identifying mitral cells is expected given a bias in viral labeling of mostly mitral cells in our experiments. Such a bias may arise from our injections aimed to target MCL, potential tropism of the Sindbis virus and/or potential bias in barcode recovery near the borders of areas during laser capture micro-dissection. However, our results show that the classifier identifies pMC with high accuracy on the BARseq data. Hence, we further applied the classifier to the (larger) MAPseq dataset (**Extended**

**Data Fig. 2d**), and found similar fractions of pMC neurons in each sampled brain (**Extended Data Fig. 2e**).

We chose the neural network parameters for the classifier using a grid search. For the grid search, we trained multiple architectures of networks multiple times (500), and further chose the architecture with lowest cross-entropy loss on average on the template dataset. The architectures tested were: single hidden layer of size 16, single hidden layer of size 20 and two hidden layers of size 10 each. These architectures were chosen to keep the number of weights low compared to the number of training data points so as to avoid overfitting. We chose to train the network on the entirety of the BARseq dataset, since our BARseq dataset is unbalanced and we had a relatively small number of data points. To check for overfitting, we compared this network to networks utilizing the same winning architecture (two hidden layers of size 10 each), but which were trained using a training and testing split approach. In particular, we split the BARseq data 90/10% and 80/20% respectively. We found that, on average, the classification performances obtained from training the classifier using the whole BARseq dataset versus training by splitting the BARseq dataset into training and testing sets were highly correlated (**Extended Data Fig. 2f, g**). Thus, training using the whole dataset appears to generalize well on this dataset. Since the result of the classification is probabilistic, we could evaluate the confidence in classification from these probabilities (**Extended Data Fig. 2h**). Consistent with robust classification of pMC neurons, most pMC (4,388 out of 4,665) were classified with more than 85% accuracy. We therefore focused subsequent analyses on this fraction of pMC neurons which was classified with high accuracy.

**Supplementary Note 3: Differences in brain-wide projections of output neurons across different domains of the olfactory bulb**

In four mice, we infected output neurons on both the dorsal and ventral surface of the olfactory bulb (**Extended Data Fig. 1a** & **3a**). Comparing the projection patterns of dorsal and ventral pMC neurons revealed small, but statistically significant differences in the distribution of projections to all six bulb target brain regions sampled (**Extended Data Fig. 3b**; $p = 0.004$ for AON, $p = 0.027$ for APC, $p = 9.9\times10^{-18}$ for PPC, $p = 0.0048$ for OT, $p = 1.2\times10^{-11}$ for CoA, and $p = 1.1\times10^{-16}$ for lENT, rank sum tests after Bonferroni correction). For example, differences observed in CoA projections were consistent with previous reports that CoA receives more input from the dorsal than ventral aspects of the olfactory bulb[17]. Importantly, the correlations between projections to different bulb target regions outlined in **Fig. 1k** were observed in both dorsal pMC and ventral pMC neurons (**Extended Data Fig. 3c**). For both the dorsal and ventral pMC neurons, we further examined whether projection strengths to particular olfactory bulb target regions were dependent on the A-P positions of neuronal somata in the bulb (**Extended Data Fig. 3d**, **e**). For projections to each bulb target brain region, we calculated the mean projection strengths of all olfactory bulb output neurons at that A-P position, relative to the mean projection strengths of shuffled neurons with randomized A-P labels (Methods). This analysis did not find statistically significant biases in projections for output neurons along the A-P axis of the olfactory bulb (**Extended Data Fig. 3d**, **e**), consistent with previous reports[17,47]. Also, similar to previous observations, we did not identify a systematic relationship between the location of bulb output neuron somata along the A-P axis of the olfactory bulb and the strength of their projections along the A-P axis of piriform cortex (**Extended Data Fig. 3f**).

**Supplementary Note 4: Correlations between individual neuronal bulb output projections along the A-P axis of piriform cortex and their co-projections to extra-piriform areas**

Mitral cell axons form orderly representations in the piriform cortex as defined by their co-projections to other brain regions. Both according to our data and previous observations[15,16], the same mitral cell usually branches and sends afferents to the piriform cortex and other, extra-piriform, brain regions. Here, for each position in the piriform cortex, we determined the conditional probability of projecting to other olfactory bulb target regions P(target|PC location). This probability describes the rate at which bulb output axons tend to connect to extra-piriform regions given that they also connect to a given position in the piriform cortex (**Fig. 2c**, **Extended Data Fig. 4b**). If the olfactory bulb-to-piriform cortex connectivity were absolutely random, these conditional probabilities would be constant throughout the piriform cortex and independent on the piriform cortex location (**Extended Data Fig. 4c**). Instead, we show that this probability displays orderly, close to linear, variation as a function of piriform cortex location (**Fig. 2c**). This implies that co-innervation of the piriform cortex and three other olfactory bulb major targets (AON, CoA and lENT) varies systematically in the piriform cortex. Our findings are therefore inconsistent with the hypothesis of random connectivity between the olfactory bulb and the piriform cortex. Interestingly, the same conditional probability of projections experiences a discontinuous change in the slope of variation at the border between the anterior and posterior piriform cortex, giving another example of the fine spatial organization of projections (**Fig. 2c**).

The spatial organization of connectivity that we report could, in principle, be understood based on the model of connection geometric locality[3,80–82]. According to this model, axons tend to form connections to nearby neurons, and the density of connections decreases as a function of distance.

Overall, however, our data suggest that a pure geometric model for spatial organization of the olfactory bulb projections cannot explain all aspects of the observed projections and instead a function-based model may be more appropriate. Several observations argue against a purely geometric explanation: 1) projections to the OT are rich and flat, despite distance to the OT varying substantially along the A-P axis of the piriform cortex; 2) the slope of spatial dependence of projection probability differs across olfactory bulb target regions (lENT vs. CoA vs. AON; **Fig. 2c**); 3) sudden changes in slope at the boundary between the APC and PPC for projections to different extra-piriform bulb target regions (**Fig. 4b**); 4) the choice of dominant projection of piriform cortex output neurons cannot be explained by proximity of the target region to the soma location within piriform cortex (e.g. projections to the olfactory bulb are stronger than those to the AON; **Fig. 3e**). We speculate that the density of connections is determined based on the functional properties of neurons in specific bulb target brain regions and the responses of specific sets of mitral cells to odorants. The geometry and locations of these regions may then have been determined based on connection length minimization *a posteriori*, in the course of evolution[80,81].

**Supplementary Note 5: Using CAV-2 retrograde labeling to validate correlations observed in the putative mitral cell projections to the lENT and PPC**

To validate some of the observations obtained via MAPseq that pMC projections to extra-piriform areas are correlated with dominant projections to specific positions along the A-P axis of piriform cortex (**Fig. 2c**), we used a CAV-2 retrograde viral labeling strategy[104] (**Extended Data Fig. 5a**). Specifically, we focused on probing the mitral cell projections to the lENT and PPC. We first injected AAV-FLEX-$\{EGFP\}_{on}$ and AAV-DIO-$\{CAR\}_{off}$ to target mitral cells on the dorsal side of OB and waited two weeks to allow optimal expression of CAR (coxsackievirus and adenovirus receptor), which acts as the receptor for CAV-2 and, thus, can help overcome potential CAV-2 tropism enabling efficient retrograde labeling. Further, we injected CAV2-Cre in either lENT or into the middle of piriform cortex (boundary between the APC and PPC) to retrogradely turn on EGFP expression in mitral cells that were also infected with AAV-FLEX-$\{EGFP\}_{on}$. Because both the lENT and the middle region of the piriform cortex have been reported to receive inputs only from mitral cells, but not tufted cells[13–16], this labeling strategy enables us to examine the distribution of mitral cell projections along the A-P axis of the piriform cortex. Consistent with our MAPseq results, CAV2-Cre injection in the lENT showed substantially stronger labeling in the posterior portion of the piriform than CAV2-Cre injection targeted to the middle of the piriform cortex (**Extended Data Fig. 5a-b**). Thus, this result further confirms our finding using MAPseq that mitral cell projections to extra-piriform projections are predictive of the distribution of their co-projections along the A-P axis of piriform cortex.

**Supplementary Note 6: Narrowly and broadly projecting putative mitral cells differentially tile the A-P axis of the piriform cortex**

Our sample of putative mitral cells (pMC) helps understand the remarkable heterogeneity in their connections. We identified two distinct classes of mitral cells according to their projection width (**Fig. 2d**, **e**). The width of projections can be characterized by the Inverse Participation Ratio (IPR), a metric describing the number of tissue slices a given cell projects to (Methods). Cells forming narrow/broad projections are described by low/high IPR respectively. Based on the sparseness of whole brain projections, pMCs from six brains were divided into two populations using a watershed algorithm (Methods): ~80% were classified as broadly projecting (BP) and the rest 20% as narrowly projecting (NP) (**Fig. 2d**, **e**, **Extended Data Fig. 6a**). Indeed, these two groups have distinct projection patterns: Broadly projecting cells project across all olfactory bulb target brain regions sampled and narrowly projecting cells mainly target the AON, APC and OT, though a subset (~50%) of them also project to PPC and CoA (**Fig. 2f**, **h**, **i**). Within each bulb target brain region, broadly and narrowly projecting mitral cells innervate distinct domains (**Extended Data Fig. 6b**): broadly projecting neurons project most strongly to the caudal portion of the anterior piriform cortex and the boundary between the anterior and posterior piriform cortex, while narrowly projecting neurons target most strongly the anterior part of APC; furthermore, within AON, broadly projecting neurons project more anteriorly than narrowly projecting neurons and more posteriorly than narrowly projecting cells to the OT, CoA and lENT. Given their different spatial distributions along the A-P axis of the piriform cortex and extra-piriform target regions, the projection gradients formed by the narrowly and broadly projecting mitral cells cannot be explained by technical issues of barcode trafficking.

We identified pMC by checking the MAPseq-based projection results against the projection patterns of mitral, tufted and deep cells found using BARseq (where cell identity is assigned based on soma location within the olfactory bulb). While the narrowly and broadly projecting cells appear to represent two subpopulations of mitral cells which differ in their projection patterns, we cannot rule out the possibility that narrowly projecting cells also include a minority of internal tufted cells displaced in the mitral cell layer (MCL)[111,112]. The projections of NP cells, however, are distinct from those of tufted cells identified by previous tracing studies[13,14] and also from those of neurons classified here as tufted cells based on their cell body location in the external plexiform layer (EPL) (**Extended Data Fig. 4a**), and include axonal branches that target, for example, the PPC and CoA. Importantly, independently of cell type identity conventions, both narrowly and broadly cell populations tile the piriform cortex in a non-random and reproducible manner and display characteristic co-innervation of specific extra-piriform bulb targets.

Interestingly, both narrowly and broadly projecting cells' axons tile (cover completely) the A-P axis of anterior piriform cortex. The tiling, however, is different for the narrowly and broadly projecting neurons. Narrowly/broadly projecting cells prefer anterior/posterior APC and follow exponential and inverted exponential distributions respectively, referenced to the anterior boundary of the piriform cortex [Eq. (1) and Eq. (2)]. While only a fraction of the sampled narrowly projecting neurons tile PPC, BP neurons projection maxima cover the full extent of the PPC following an exponential distribution [Eq. (1)]. The tiling by narrowly projecting cells reveals a potential pitfall for tracing studies involving multiple neurons at the same time. When multiple narrowly projecting cells are traced, they may appear to form a broad projection spanning the entire A-P axis of the anterior piriform cortex, while individual neurons form specific projections, even if these cells may originate in a single glomerulus[15,16]. Thus, when examining the specificity of the olfactory bulb-to-piriform cortex connectivity, both single-neuron resolution and high throughput are necessary.

Are projections of the same olfactory bulb output neuron to the anterior and posterior piriform cortex correlated? To answer this question, we examined the simultaneous tiling of these piriform cortex subdivisions by projection peaks of the same neurons. The same NP neuron can form projections in both the APC and PPC and locations of the peaks of in projection can be identified in each piriform subdivision (**Fig. 2h**). As discussed above, narrowly projecting cells (**Fig. 2h**) display exponential tiling [Eq. (1)] along the A-P axis of the anterior piriform cortex. The same is true for the narrowly projecting cells which target the posterior piriform cortex. When the same neurons are re-sorted according to the location of PPC projection maxima (**Fig. 2i**), they follow exponential tiling similar to APC. For the same neuron, the locations of peaks in the anterior and posterior piriform cortex are however uncorrelated (**Fig. 2j**). A similar feature is observed in the tiling distributions by broadly projecting neurons. For the same neuron, locations of peaks in projection density in the anterior and posterior piriform cortex appear to be independent (**Fig. 2k, l**) with no observed correlation (**Fig. 2m**). Because broadly projecting cells display an inverted exponential tiling in the anterior piriform cortex [Eq. (2)], these neurons form denser projections in the posterior APC and anterior PPC respectively, thus contributing to the peak in projection density observed at the boundary between the anterior and posterior piriform cortex (**Fig. 2m**). Overall, our findings suggest that connectivity of the same bulb output cell to the APC and PPC is not correlated. Within a scenario in which the A-P locations of projections in the anterior and posterior piriform cortex correlate with the representation of two variables important for olfactory processing, these variables appear independent. Further investigation is necessary to determine the functional implications of the piriform cortex tiling by bulb projections and of the strong innervation of the APC / PPC boundary region suggestive of anatomical modules beyond the APC and PPC cortical subdivisions.

**Supplementary Notes 7: Intra-piriform and brain-wide piriform cortex projections and comparisons with previous work**

MAPseq analysis of intra-piriform connectivity and brain-wide organization of piriform cortex output neurons recapitulate previous observations[49,84–87,92]. First, clustering of piriform cortex slices based on their bi-directional intra-piriform connectivity across cortical slices identified two groups of slices that were highly connected within each group, but sparsely connected across groups. These two groups corresponded to the anatomically-defined APC and PPC, respectively, indicating that differences found in intra-piriform connectivity match the anatomically defined APC-PPC boundary (**Extended Data Fig. 7a**). Second, consistent with previous observations on connectivity within the piriform cortex, APC-to-PPC projections were stronger than PPC-to-APC projections (**Fig. 3b**, **c**). Third, at the population level, the projection strengths of the piriform cortex outputs to specific target regions were consistent with bulk tracing data obtained using the Allen Brain Atlas (**Extended Data Fig. 7b-d**). Furthermore, the APC-to-APC and PPC-to-PPC correlations between our data and the Allen Brain atlas were significantly higher than correlations between APC and PPC across these datasets (**Extended Data Fig. 7e**). Overall, these observations indicate that MAPseq projection data capture well previously reported APC versus PPC differences in both piriform cortex output and intra-piriform connectivity.

**Supplementary Notes 8: Piriform cortex output neuron groups differ in their dominant projection targets along the A-P axis of the cortex**

We used two layers of Louvain community detection[109] to cluster piriform cortex projection neurons into groups, then manually combined groups from the second layer that did not appear disjointed when visualized using t-SNE (**Fig. 3e**; **Extended Data Fig. 8a**). This resulted in six groups that were differentially distributed along the A-P axis of the piriform cortex (**Fig. 3f**; **Extended Data Fig. 8b**, **c**) in a manner reproducible across brains (**Extended Data Fig. 8c**). The projection patterns of these piriform cortex output neurons partially predicted the A-P locations of their somata compared to the full output projection patterns (Methods). Prediction errors were 1.0 ± 0.1 mm (mean ± sem) when using the six groups identity only, compared to 1.3 ± 1.4 mm when using shuffled data, and 0.7 ± 0.5 mm when using the full projection patterns of the sampled piriform cortex output neurons (**Extended Data Fig. 8d**, **e**). Interestingly, additional, finer gradients of projection along the A-P axis were observed within some of the groups (**Fig. 3e**). For example, while neurons in group 5 were heterogeneous in projecting to many piriform cortex targets, their distribution of along the A-P axis of piriform cortex showed a bimodal projection distribution that correlated specifically with projections to the striatum and the contralateral piriform cortex: neurons in the anterior portion of piriform cortex mostly projected to caudate-putamen (CP), whereas neurons in the posterior portion of piriform cortex also projected to nucleus accumbens (ACB) and the contralateral piriform cortex.

Importantly, the groups of piriform cortex neurons defined by their output projection targets also showed different biases in their intra-piriform projections: e.g., neurons in group 1, which projected to AON, were more likely to project anterior within the piriform cortex ($p = 3.6 \times 10^{-94}$, Sign-rank test after Bonferroni correction), whereas neurons in group 3, which projected to lENT ($p = 1.9 \times 10^{-303}$, sign-rank test after Bonferroni correction), were more likely to project posterior within the piriform cortex (**Extended Data Fig. 8f**). We cannot rule out, however, the possibility that some of these differences in intra-piriform projections were contributed by passing fibers of the output projections.

Taken together, these findings reveal the existence of systematic gradients, reproducible across individuals, along the A-P axis of the piriform cortex, both in terms of projection modules within the piriform cortex, as well as with respect to the organization of piriform cortex outputs to other brain regions.

**Supplementary References:**

110. Fürth, D. *et al.* An interactive framework for whole-brain maps at cellular resolution. *Nat. Neurosci.* **21**, 139–149 (2018).

111. Schwarz, D. *et al.* Architecture of a mammalian glomerular domain revealed by novel volume electroporation using nanoengineered microelectrodes. *Nat. Commun.* **9**, 1–14 (2018).

112. Mori, K., Kishi, K. & Ojima, H. Distribution of dendrites of mitral, displaced mitral, tufted, and granule cells in the rabbit olfactory bulb. *J. Comp. Neurol.* **219**, 339–355 (1983).

#### Extended Data Figure 1

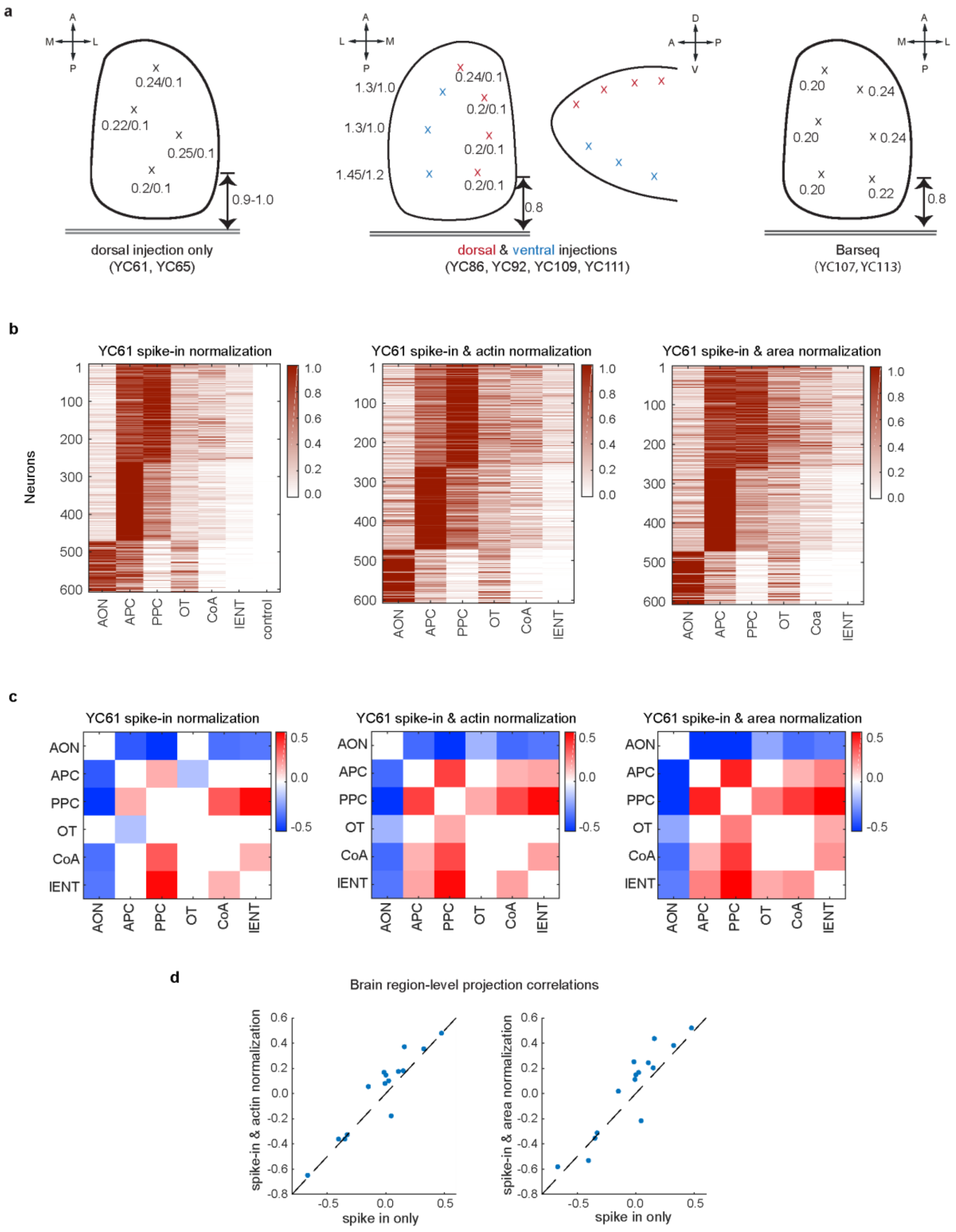

**Extended Data Figure 1. Olfactory bulb barcoded Sindbis virus injections and normalization schemes**

*(**a**) Schematics for multiple site injections in the olfactory bulb of the barcoded Sindbis virus. The most posterior injection sites are 0.8-1.0 mm rostral to the major blood vessel behind the bulb. Brains of YC61&YC65 (left) were injected at four sites (indicated by x) at two depth of 0.2-0.25 mm and 0.1 mm (indicated next to injection site) on dorsal side of the bulb. YC86, YC92, YC109, YC111 (middle) were injected at multiple sites on both dorsal (indicated as red x) and ventral aspects of the bulb (indicated by blue x). Each site was injected at two depths (distance in mm next to injection site in schematics). BARseq Brains YC107&YC113 (right) were injected at six sites (indicated as x) on the dorsal OB at a single depth of 0.2-0.25 mm (indicated next to injection site). Details of injection coordinates are provided in Methods. (**b**) Brain area resolution projection matrices of one example brain (YC61) normalized based on spike-in RNA only, spike-in RNA and β-actin (measured by qPCR), and spike-in RNA and dissected brain area size (Methods). Projection strength of each barcode has been normalized to the maximum projection across different brain regions. Data have been arranged into three groups identified by Louvain Community Detection. (**c-d**) Pearson correlation between olfactory bulb projections to different brain regions for data normalized to spike-in RNA only, spike-in RNA and actin and spike-in RNA and dissected brain area size for one example data set (YC61). Only statistically significant correlations (after Bonferroni correction) are shown. Each dot in panel (**d**) represents a Pearson correlation between bulb projections to different brain regions for data normalized to spike in RNA only (x-axis) and spike-in RNA and actin (y-axis, left) or spike-in RNA and dissected brain area size (y axis, right).*

#### Extended Data Figure 2

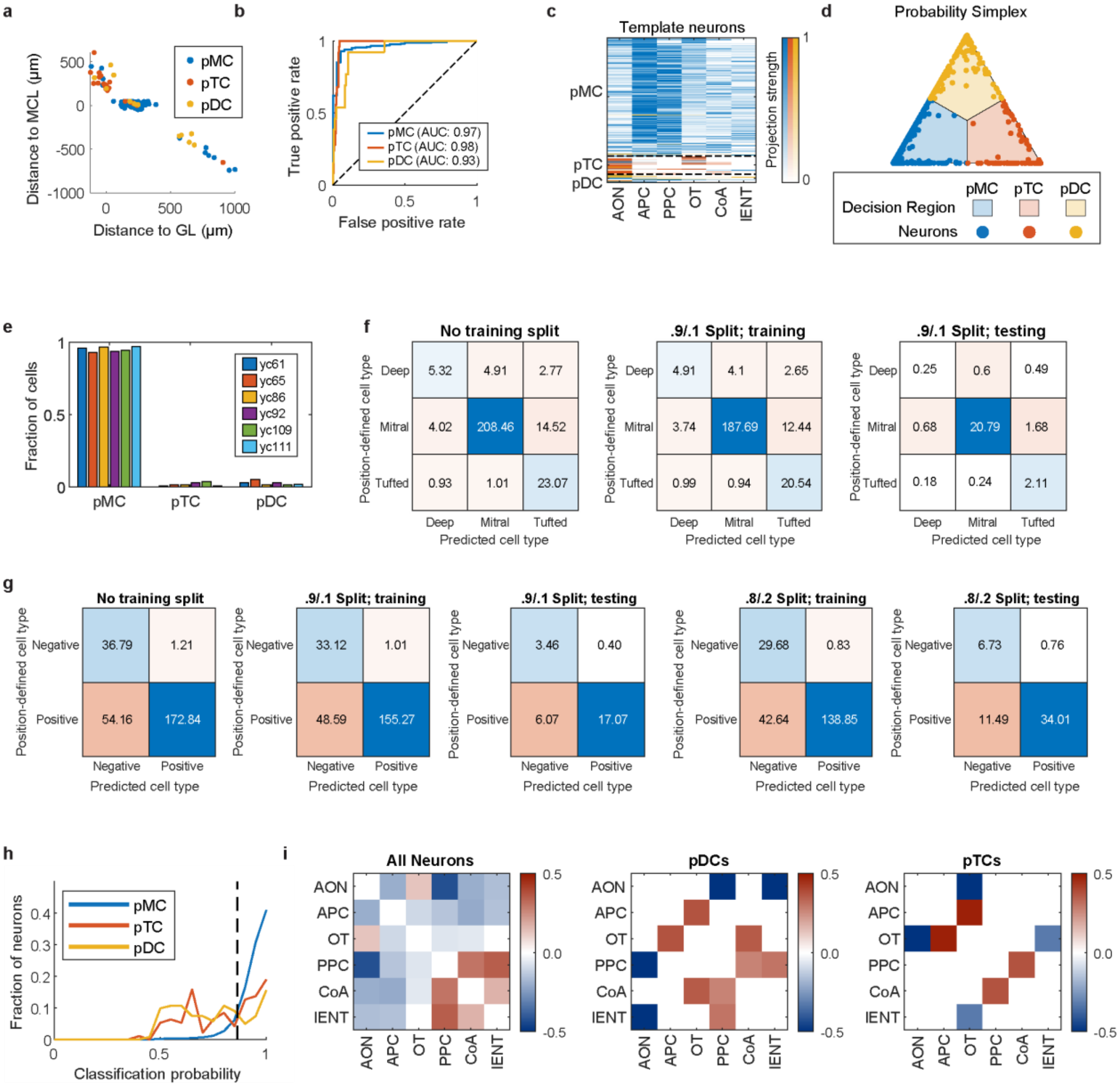

**Extended Data Figure 2. Classification of the olfactory bulb output neurons based on their projection patterns using BARseq data as template.**

*(**a**) Template neurons identified via BARseq shown in physical space (distance to mitral cell layer, MCL vs. distance to glomerular layer), color coded by the classifier assigned identities (blue – pMC, red – pTC and yellow - pDC). (**b**) Receiver Operating Characteristic (ROC) analysis of the classifier for all three classes. (**c**) Projection patterns of the template neurons. Neurons are sorted according to classifier results and color coded by their true labels based on soma postions in the olfactory bulb (blue - pMC, red - pTC, yellow - pDC). (**d**) MAPseq neurons shown in the space of their classification probabilites, projected onto a 2d plane. (**e**) Fraction of classes identified in each individual brain (color coded). (**f**) The classification confusion matrix of template neurons, determined by the maximum probability of classification. Matrix values shown were averaged over 500 trained networks, trained on the entire dataset (left), and using a 90%/10% training (center) and testing (right) split. (**g**) The classification confusion matrix of template neurons, using a thresholding criterion of pMCs probability larger than 85%. Matrix values shown were averaged over 500 trained networks, trained on the entire dataset (left), using a 90%/10% training/testing (center-left/center) split, and a 80%/20% training/testing (center-right / right) split. (**h**) The distribution of classification probability for each class. Dashed vertical line indicates 85% probability threshold. (**i**) Pearson correlation between projections to different areas in all neurons, pDC, and pTC. Only statistically significant correlations (after Bonferroni correction) are shown.*

#### Extended Data Figure 3

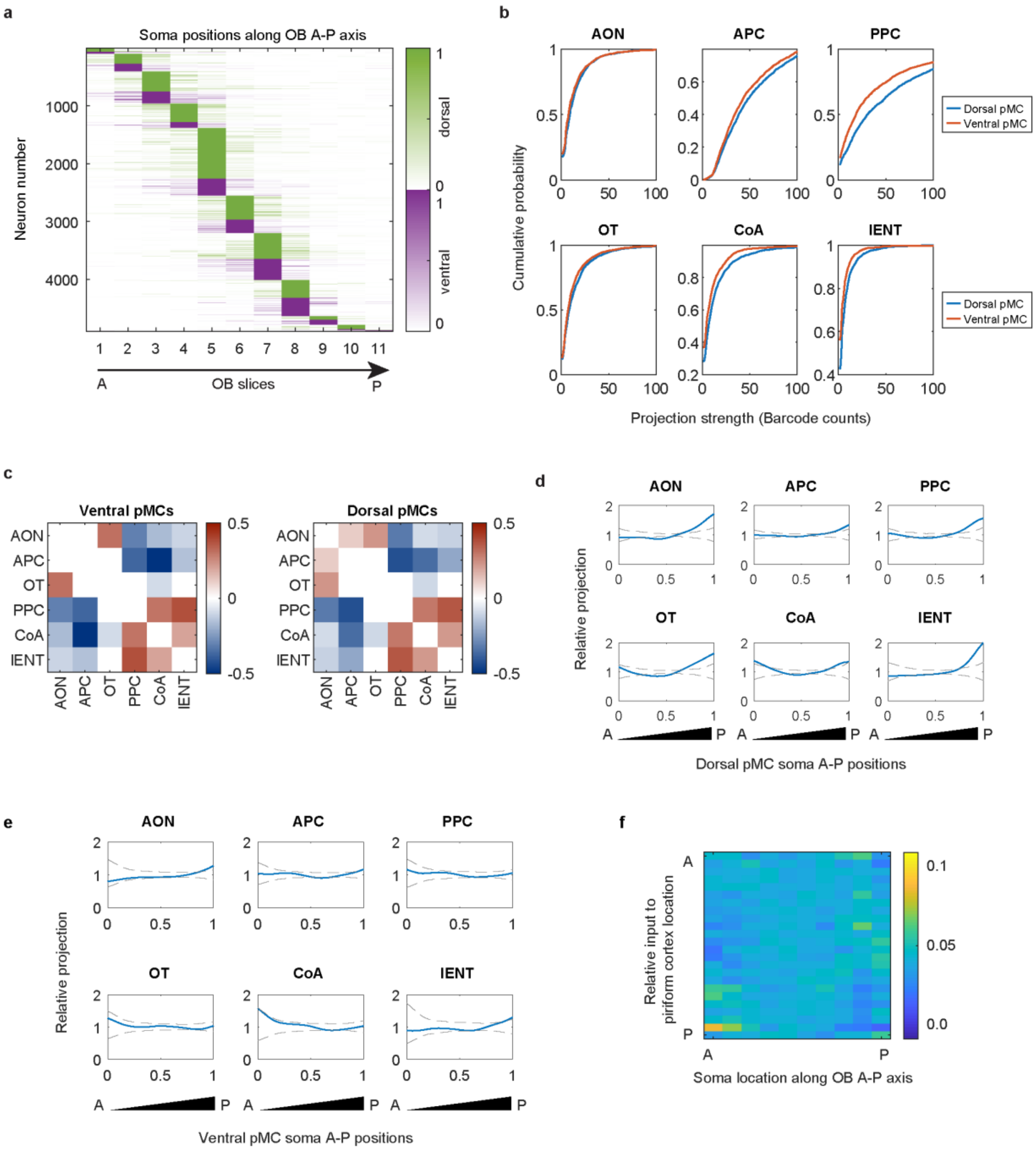

**Extended Data Figure 3. Spatial distribution of barcode labeled pMC somata and projection patterns as a function of different locations within the olfactory bulb**

*(**a**) Barcode counts in the olfactory bulb. Neurons are sorted by the peak positions, which indicate the soma locations along the A-P axis of OB. Projection patterns are color coded by the dorsal (green) and ventral (purple) location of their somas. (**b**) Cumulative probability distribution of the strength of projections of dorsal and ventral pMC to the indicated brain regions. (**c**) Pearson correlation between projections to different bulb target regions for the pMC neurons from the ventral and dorsal aspects of the bulb. Only statistically significant projections are shown. (**d-e**) Relative projection strengths of dorsal (**d**) and ventral (**e**) pMC from specific A-P positions in the bulb (x-axis) to the indicated olfactory bulb target regions. The projection strength was normalized to the mean projection strength of all dorsal or ventral pMC neurons. Dashed lines indicate 95% confidence interval obtained by shuffling the A-P positions of neurons. (**f**) Relative projection strengths of pMC neurons to different piriform cortex A-P positions for neurons at different A-P locations within the olfactory bulb. Each column shows the ratio between the mean projection strengths of neurons at the indicated A-P locations in the olfactory bulb (x-axis) to different A-P locations in the piriform cortex (y-axis) and the mean projection strengths of all neurons to different A-P locations in the piriform cortex (y-axis). These ratios are further normalized by the sum of each column to allow visualization.*

#### Extended Data Figure 4

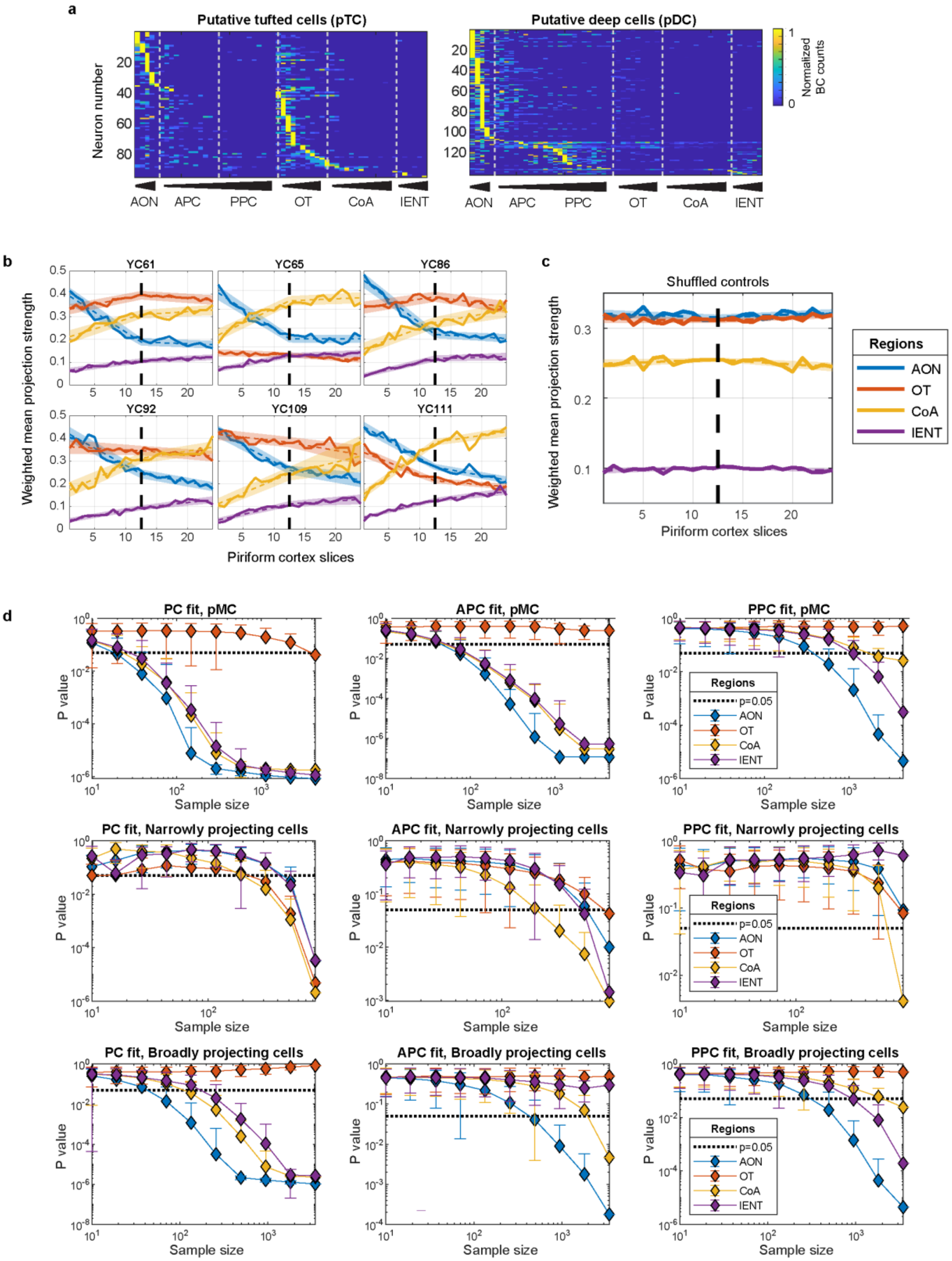

**Extended Data Figure 4. Reproducibility across individuals, shuffled controls and downsampling analysis on the projection patterns of the olfactory bulb output neurons**

(***a***) *Slice-level (200 µm resolution) projections of pTC and pDC neurons, sorted by the location of their peak positions along the A-P axis of each of the olfactory bulb target regions sampled. Projections were normalized to the maximum projection within each row. (**b-c**) Weighted mean co-innervation strength across piriform cortex slices and extra-piriform areas [the conditional probability of co-innervation P(target|PC location), solid lines] in each of the six brains sampled (**b**) and the same data obtained by shuffling the piriform cortex slices (**c**). Dashed lines/shaded areas show piecewise linear fits in APC and PPC with the 95% confidence interval obtained by bootstrap. (**d**) Distribution of p-values of Spearman correlations after down sampling. The Spearman correlations are calculated between the A-P position of the piriform cortex slice and the weighted mean co-innervation strengths between target regions and the piriform cortex slice considered. Spearman correlation was calculated across the entire piriform cortex (left column), only on APC (center column) and only on PPC (right column). Barcodes used for down sampling were selected among all pMC neurons (top row), narrowly projecting neurons (center row) and broadly projecting neurons (bottom row). Sample sizes are as shown on the x-axis of each plot.*

#### Extended Data Figure 5

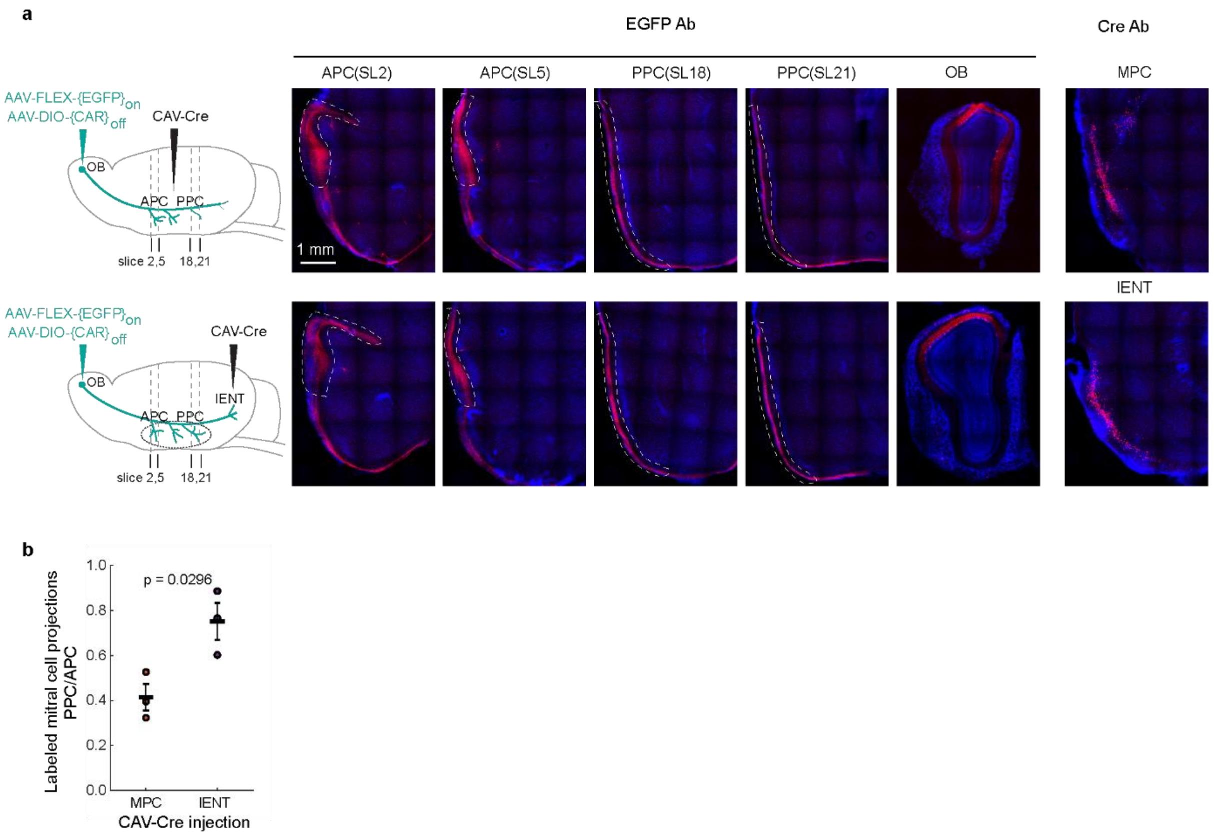


**Extended Data Figure 5. Validation of the spatial organization of pMC projections using CAV-Cre bulk retrograde labeling**

*(**a**) Left, schematics of CAV-Cre retrograde labeling in the middle of piriform cortex (MPC, top panel) and lENT (bottom panel). Center, immunofluorescence of labeled mitral cells at different A-P positions of piriform cortex and olfactory bulb. Mitral cell projections to piriform cortex are largely restricted to layer I, as indicated by white dashed lines. Right, immunofluorescence of Cre at MPC (top panel) and lENT (bottom panel). (**b**) The ratio of projection strength of labeled mitral cells (as measured by EGFP immunofluorescence) in the posterior portion of the piriform cortex (slices 17-22) to the labeled mitral cell projection strength in the anterior portion of the piriform*

*cortex (slices 1-6). Each dot represents data collected from an animal. Bars indicate mean ± SEM. n=3 mice in each group, p=0.03 by Student's t test.*

#### Extended Data Figure 6

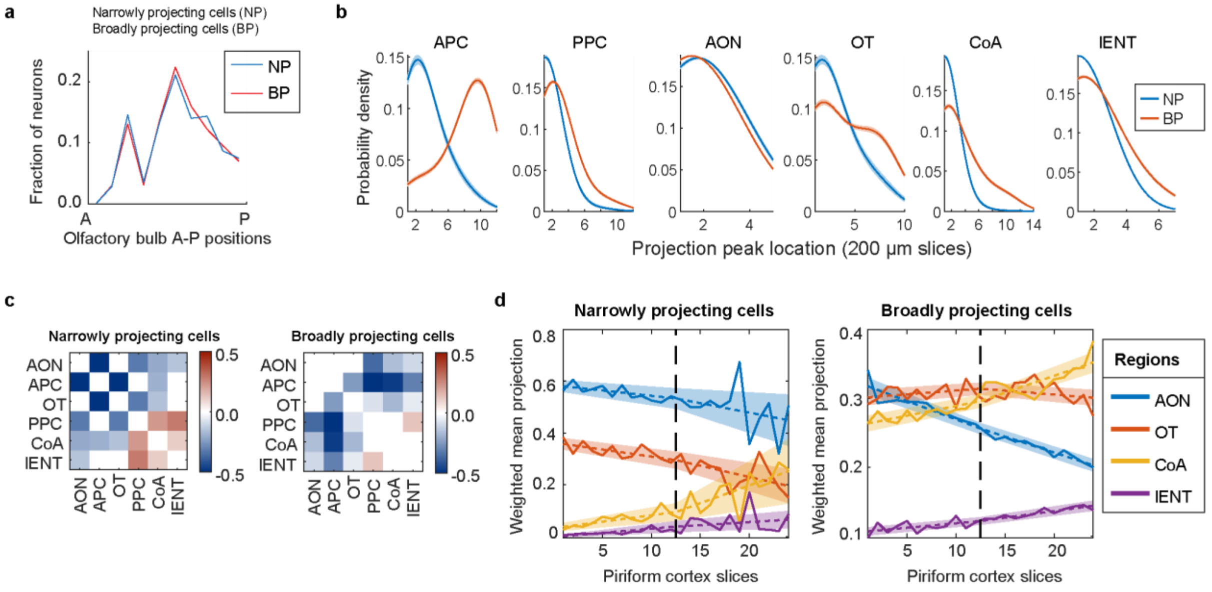


**Extended Data Figure 6. Spatial organization of the projections of broadly projecting and narrowly projecting cells**

*(**a**) The fraction of broadly and narrowly projecting neurons' somata along the A-P axis of the olfactory bulb. (**b**) Distribution of projection peak positions, smoothed using a gaussian kernel, in the indicated areas for the narrowly and broadly projecting putative mitral cells. (**c**) Pearson correlation between projections to different areas of the narrowly and broadly projecting neurons. Only statistically significant (after Bonferroni correction) correlations are shown. (**d**) Mean projection strengths (solid lines) in the indicated extra-piriform bulb target brain regions weighted by projection strengths to the indicated A-P position in the piriform cortex (x-axis) for the broadly and narrowly projecting neurons. Dashed lines indicate piecewise linear fits in APC and PPC and shaded areas indicate range of fits from bootstrapping.*

Extended Data Figure 7

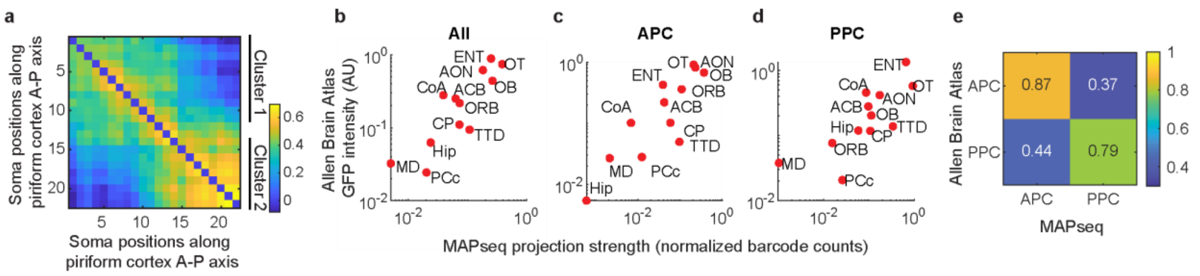


**Extended Data Figure 7. Intra-piriform and brain-wide projections of piriform cortex output neurons referenced to the Allen Brain Atlas**

*(**a**) Spearman correlation between barcode counts in different piriform cortex slices. Clusters indicated on the right indicate highly inter-connected slices obtained by Louvain community detection (see Methods). (**b-d**) Projection strengths from the Allen Brain Atlas (ABA) (y-axis) and sum of barcodes of neurons from MAPseq (x-axis) for the indicated areas for all the piriform cortex neurons (**b**), APC neurons (**c**), and PPC neurons (**d**). (**e**) Pearson correlations between the projection strengths from the Allen Brain Atlas (ABA) and barcode counts from MAPseq separately for APC and PPC.*

Extended Data Figure 8

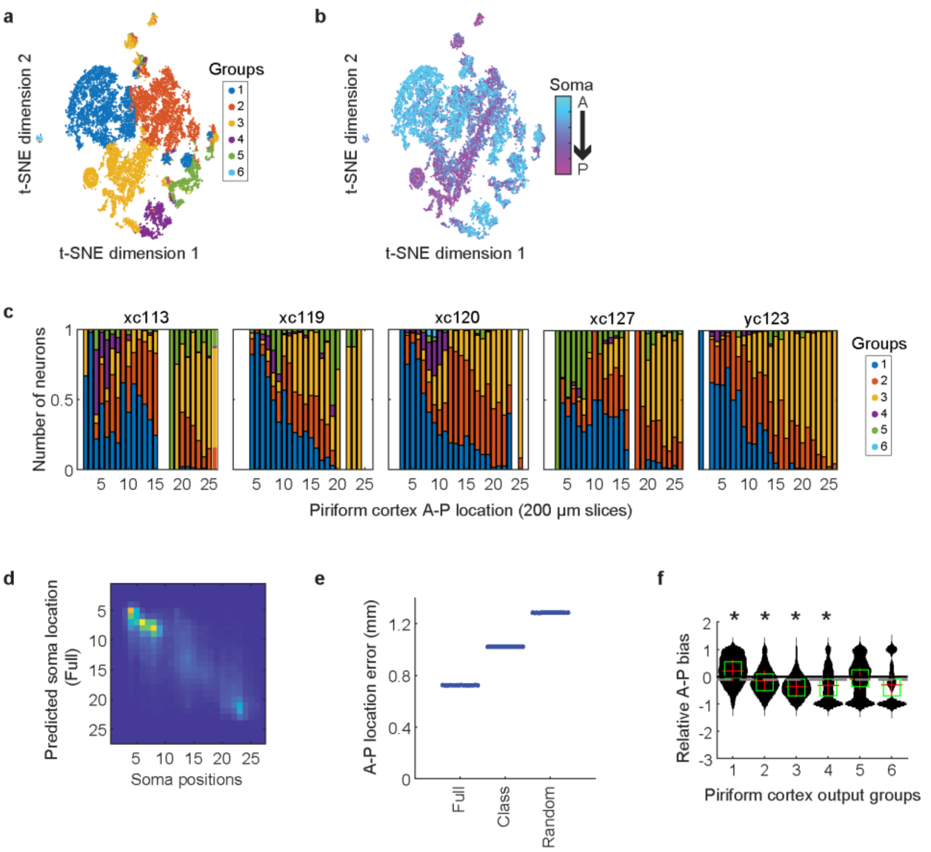


**Extended Data Figure 8. Differential brain-wide projections of groups of piriform cortex output neurons organized in a graded manner along the A-P axis.** *(**a**-**b**) t-SNE plots of the piriform cortex output projections color coded by groups defined by the output projections (**a**) and soma locations (**b**). (**c**) The fraction of neurons belonging to each group at the indicated A-P positions, plotted separately for each brain. (**d**) Prediction of soma locations (y-axis) using full projection pattern. (**e**) Mean prediction errors of the A-P positions of piriform cortex neuronal somata using full projection pattern, group identities, and randomized output projection patterns. Individual dots indicate trials using cross validation. (**f**) Differences in intra-piriform cortex projections in the anterior and posterior directions for each piriform cortex projection group identified (Methods). Positive numbers indicate stronger anterior projections and negative*

*numbers stronger posterior projections. Dashed line indicates mean projection bias for all the piriform cortex output neurons; * $p < 10^{-10}$ compared to no bias using sign rank tests after Bonferroni correction.*

#### Extended Data Figure 9

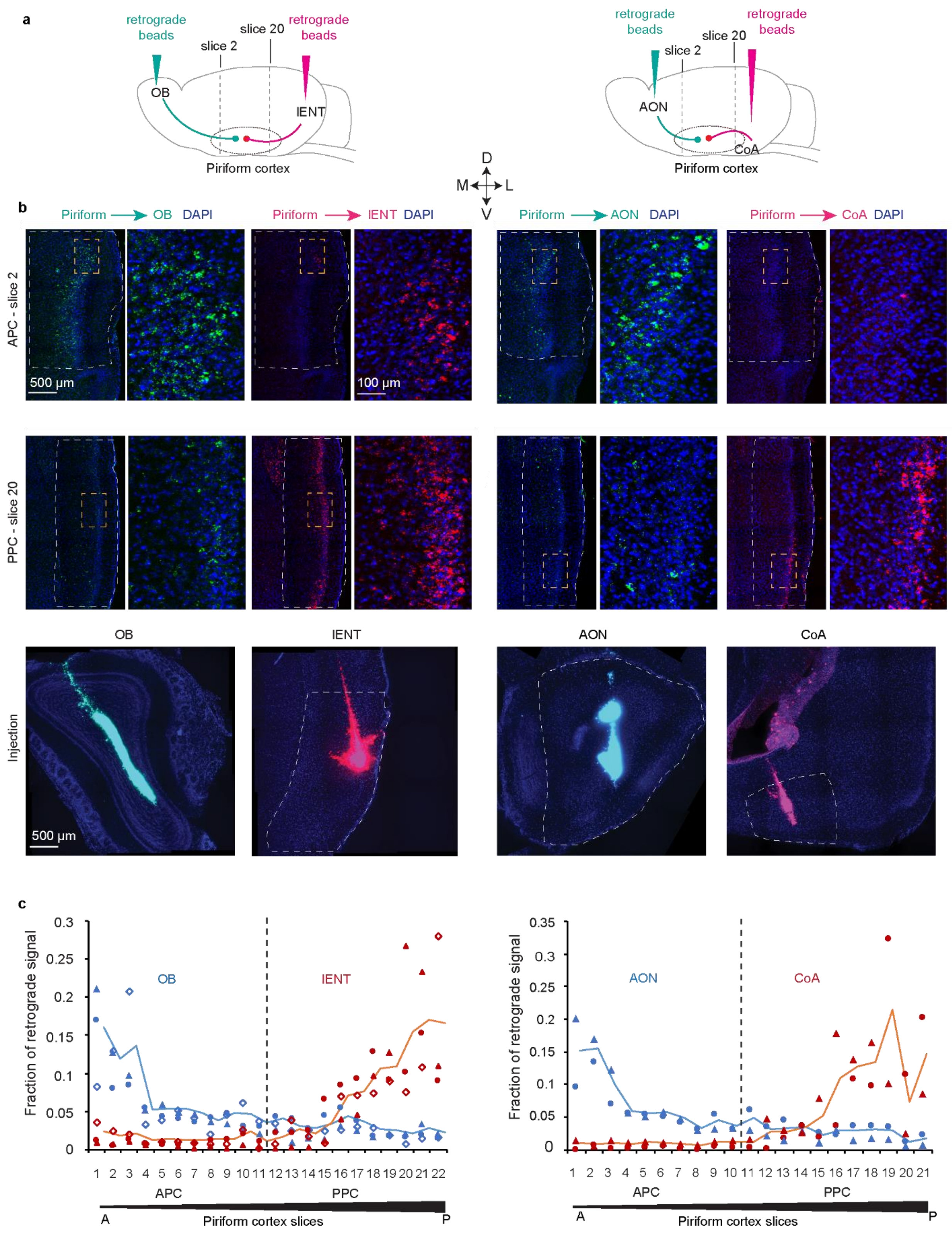

**Extended Data Figure 9. Validation of piriform cortex output projection patterns along A-P axis via bulk retrograde tracing.**

*(**a**) Cartoon schematics of injecting retrograde beads of different colors into the olfactory bulb vs. lENT and the AON vs. CoA. (**b**) Representative example images of retrograde labeling at different A-P positions in the piriform cortex and labeling at different injection sites. A selected area (golden rectangle) in each piriform cortex slice was amplified on the right. (**c**) Quantification of retrograde labeling from different brain regions along the A-P axis of the piriform cortex. Retrograde fluorescence signals in the piriform cortex from bulb injections (Left) and AON injections (Right) are shown in blue; retrograde signals from lENT (Left) and CoA (Right) are shown in red. Different shapes represent different animals (3 mice for co-injection into OB & lENT; 2 mice for co-injection into AON & CoA); solid lines indicate the mean across different animals. Individual piriform cortex slices are 100 µm coronal sections arranged from the anterior to the posterior end of the cortex and spaced 200 µm apart.*

**Supplementary Table 1**. Animals used in the study

| # of animals | Strain | Experiment | Injection | Processing Conditions | Injection Date | ID | MAPseq library |
|---|---|---|---|---|---|---|---|
| 2 | C57BL/6J, 8-10-week old, male, Jackson Laboratory | OB MAPseq | 50 nl Sindbis Viral library (1:3 diluted). Two depths of 100 µm and 200-250 µm (from brain surface) at four penetration sites along the A-P axis of the dorsal aspect of the bulb (0.8 mm, 1.2 mm, 1.7 mm, 2.2 mm anterior to the blood vessel between OB and prefrontal cortex, 0.7-1.0 mm lateral from midline) | 44 hrs post injection, fresh frozen for MAPseq | 5/4/2018<br>6/8/2018 | YC61<br>YC65 | ZL153<br>ZL153 |
| 4 | C57BL/6J, 8-10-week old, male, Jackson Laboratory | OB MAPseq | 50 nl Sindbis Viral library (1:3 diluted). Two depths of 100 µm and 200-250 µm (from brain surface) at four penetration sites along the A-P axis of the dorsal aspect of the bulb (0.8 mm, 1.2 mm, 1.7 mm, 2.2 mm anterior to the blood vessel between OB and prefrontal cortex, 0.7-1.0 mm lateral from midline).<br>And Two depths at 3 penetration sites on ventral aspect of the bulb (0.75 mm anterior, 1.2-1.1 mm lateral, 1.2 and 1.45 mm deep; 1.3 mm anterior, 1.0-1.1 mm lateral, 1.0 and 1.3 mm deep; 1.8mm anterior, 0.8-0.9mm lateral, 1.0 and 1.3mm deep) | 44 hrs post injection, fresh frozen for MAPseq | 1/23/2019<br>2/28/2019<br>8/7/2019<br>8/29/2019 | YC86<br>YC92<br>YC109<br>YC111 | ZL172<br>ZL172<br>ZL196<br>ZL196 |
| 1 | C57BL/6J, 9 weeks old, male, Jackson Laboratory | OB BARseq | 70 nl Sindbis Viral library (1:3 diluted). One depth of 200-250 µm from brain surface into six sites, spaced 500 µm apart along the A-P axis of the dorsal aspect of the bulb. | 24 hrs post injection, PFA fixation of OB for 24 hrs, cryoprotection, and in situ sequencing. Fresh frozen the the rest of the brain (apart from OB) for MAPseq | 7/17/2019 | YC107 | ZL212 |
| 1 | C57BL/6J, 8 weeks old, male, Jackson Laboratory | OB BARseq | 70 nl Sindbis Viral library (1:3 diluted). One depth of 200-250 µm from brain surface into six sites, spaced 500 µm apart along the A-P axis of the dorsal aspect of the bulb | 24 hrs post injection, fresh frozen for in situ sequencig and MAPseq | 12/1/2019 | YC113 | ZL212 |

| | | | | | | | |
|---|---|---|---|---|---|---|---|
| 5 | C57BL/6J, 8-10-week old, male, Jackson Laboratory | PC MAPseq | 100 nl Sindbis Viral library (1:3 diluted). Three sites along the A-P axis of piriform cortex (anterior: AP +1.75 mm, ML 2.8 mm, DV 4.75 mm from the skull surface of bregma; middle: AP +0.15 mm, ML 3.9 mm, DV 5 mm from the skull surface of bregma; posterior: AP -1.5 mm, ML 4.25 mm, DV 5.5 mm from the skull surface of bregma) | 24 hrs post injection, 4% PFA perfusion, 24 hrs postfix, MAPseq | 11/7/2019<br>12/9/2019<br>12/9/2019<br>1/15/2020<br>2/18/2020 | XC113<br>XC119<br>XC120<br>XC127<br>YC123 | ZL206<br>ZL208(targets & injection)&ZL213 (injection)<br>ZL208(targets & injection)&ZL214 (injection)<br>ZL222(targets & injection),<br>ZL227(targets)<br>ZL228(targets)&ZL229(injetion) |
| 3 | C57BL/6J, 6-7-week old, male, Jackson Laboratory | OB-MPC Retrograde CAV2-Cre | 70 nl viral mixture of AAV pCAG-FLEX-{EGFP}on (Addgene plasmid #51502, titer 1.1E+12 GC/ml,) and AAV-{CAR}off ( titer 1E+12 GC/ml) into 200 µm depth from the bulb surface at three penetration sites along the A-P axis of the OB dorsal aspect. 2 weeks later, 70nl of CAV2-Cre (Montpellier vectorology platform, titer 2.75E+12 pp/ml) into each site of MPC: AP 0.10 mm, ML 3.5 mm, DV 5.5 mm and AP 0.10 mm, ML 3.85 mm, DV 5 mm from the skull surface of bregma. | 2 weeks post CAV2-Cre injection, 4% PFA perfusion, 24 hrs postfix, immunostaining for EGFP signals. | 7/22&8/7/2020<br>9/1&9/15/2020<br>11/26&12/11/2020 | YC145<br>YC170<br>YC182 | N/A |
| 3 | C57BL/6J, 6-7-week old, male, Jackson Laboratory | OB-ENT Retrograde CAV2-Cre | 70 nl viral mixture of AAV pCAG-FLEX-{EGFP}on (Addgene plasmid #51502, titer 1.1E+12 GC/ml,) and AAV-{CAR}off ( titer 1E+12 GC/ml) into 200µm depth from the bulb surface at three penetration sites along the A-P axis of the OB dorsal aspect. 2 weeks later, 140nl of CAV2-Cre (Montpellier vectorology platform, titer 2.75E+12 pp/ml) into lENT: AP -3.5 mm, ML 4.4 mm, DV 4.7 mm from the skull surface of bregma. | 2 weeks post CAV2-Cre injection, 4% PFA perfusion, 24 hrs postfix, immunostaining for EGFP signals. | 5/29&6/15/2020<br>7/22&8/7/2020<br>9/1&9/15/2020 | YC132<br>YC146<br>YC150 | N/A |

| | | | | | | | |
|---|---|---|---|---|---|---|---|
| 3 | C57BL/6J, 8-10-week old, male, Jackson Laboratory | OB vs ENT Retrograde microbeads | 35 nl different fluorescent microbeads (full strength green beads and 1:4 diluted read beads). OB: 1.2 mm anterior to the blood vessel between OB and prefrontal cortex, 0.9 mm from the middle line, 0.6 and 1.25 mm deep from the bulb surface; lENT: AP -3.5 mm, ML 4.5 mm, DV 4.75 mm from the skull surface of bregma | 3-4 days post injection, 4% PFA perfusion, 24 hrs postfix | 5/28/2020<br>6/4/2020<br>7/15/2020 | YC128<br>YC129<br>YC140 | N/A |
| 2 | C57BL/6J, 8-10-week old, male, Jackson Laboratory | AON vs CoA Retrograde microbeads | 35 nl different fluorescent microbeads (full strength green beads and 1:4 diluted read beads). AON: AP 3.2 mm, ML1.25 mm, DV 3.8 mm from the skull surface of bregma; CoA: AP -2 mm, ML2.5 mm, DV 5.8 mm from the skull surface of bregma. | 3-4 days post injection, 4% PFA perfusion, 24 hrs postfix | 6/19/2020<br>7/3/2020 | YC135<br>YC137 | N/A |

**Supplementary Table 2**. Filters and lasers used for histology

| Channel | Laser (nm) | Excitation | Dichroic | Emission (Filter or collection wavelength) |
|---|---|---|---|---|
| DAPI | 405 | zet402/468/555/640x (Chroma) | Zt402/468/555/640rpc-ufs (Chroma) | Zet402/648/555/640m (Chroma) |
| G/YFP | 520 | Zet443-518x (Chroma) | Zt443-518rpc (Chroma) | FF01-565/24 (Semrock) |
| T/RFP | 555 | zet402/468/555/640x (Chroma) | Zt402/468/555/640rpc-ufs (Chroma) | FF01-585/11 (Semrock) |
| A/Cy5 | 640 | zet402/468/555/640x (Chroma) | FF652-Di01 (Semrock) | FF01-676/29 (Semrock) |
| C | 640 | zet402/468/555/640x (Chroma) | FF652-Di01 (Semrock) | FF01-725/40 (Semrock) |
| GFP | 470 | zet402/468/555/640x (Chroma) | Zt402/468/555/640rpc-ufs (Chroma) | FF01-525/30 (Semrock) |
| TexasRed | 555 | zet402/468/555/640x (Chroma) | Di02-R594 (Semrock) | FF01-647/57 (Semrock) |

**Supplementary Table 3**. Summary statistics of figures. (See "Supp table 3.xlsx")

**Supplementary Table 4**. Piriform cortex target regions sampled via MAPseq

| **Brain region name abbreviation** | **Brain region name** | **Allen Atlas coronal levels** |
|---|---|---|
| OB | Olfactory bulb | 1-20 |
| OBc | contralateral olfactory bulb | 1-10 |
| AON | Anterior olfactory nucleus | 22-28 |
| ORB | Orbital area and agranular insular area | 24-33 |
| TTD | Taenia tecta, dorsal part and lateral septal nucleus, rostroventral part | 34-46 |
| OT | Olfactory tubercle | 34-53 |
| ACB | Nucleus accumbens | 40-50 |
| CP | Caudoputamen | 40-50 |
| APCc | contralateral anterior piriform cortex | 32-53 |
| PPCc | contralateral posterior piriform cortex | 54-79 |
| CoA | Cortical amygdala | 66-82 |
| TH | Mediodorsal nucleus of the thalamus | 64-74 |
| Hip | Hippocampus | 78-90 |
| lENT | lateral entorhinal cortex | 82-100 |
